# *Staphylococcus aureus* counters organic acid anion-mediated inhibition of peptidoglycan cross-linking through robust alanine racemase activity

**DOI:** 10.1101/2024.01.15.575639

**Authors:** Sasmita Panda, Yahani P. Jayasinghe, Dhananjay D. Shinde, Emilio Bueno, Amanda Stastny, Blake P. Bertrand, Sujata S. Chaudhari, Tammy Kielian, Felipe Cava, Donald R. Ronning, Vinai C. Thomas

## Abstract

Weak organic acids are commonly found in host niches colonized by bacteria, and they can inhibit bacterial growth as the environment becomes acidic. This inhibition is often attributed to the toxicity resulting from the accumulation of high concentrations of organic anions in the cytosol, which disrupts cellular homeostasis. However, the precise cellular targets that organic anions poison and the mechanisms used to counter organic anion intoxication in bacteria have not been elucidated. Here, we utilize acetic acid, a weak organic acid abundantly found in the gut to investigate its impact on the growth of *Staphylococcus aureus*. We demonstrate that acetate anions bind to and inhibit D-alanyl-D-alanine ligase (Ddl) activity in *S. aureus*. Ddl inhibition reduces intracellular D-alanyl-D-alanine (D-Ala-D-Ala) levels, compromising staphylococcal peptidoglycan cross-linking and cell wall integrity. To overcome the effects of acetate-mediated Ddl inhibition, *S. aureus* maintains a substantial intracellular D-Ala pool through alanine racemase (Alr1) activity and additionally limits the flux of D-Ala to D-glutamate by controlling D-alanine aminotransferase (Dat) activity. Surprisingly, the *modus operandi* of acetate intoxication in *S. aureus* is common to multiple biologically relevant weak organic acids indicating that Ddl is a conserved target of small organic anions. These findings suggest that *S. aureus* may have evolved to maintain high intracellular D-Ala concentrations, partly to counter organic anion intoxication.

**Significance:** Under mildly acidic conditions, weak organic acids like acetic acid accumulate to high concentrations within the cytosol as organic anions. However, the physiological consequence of organic anion accumulation is poorly defined. Here we investigate how the acetate anion impacts *S. aureus*. We show that acetate anions directly bind Ddl and inhibit its activity. The resulting decrease in intracellular D-Ala-D-Ala pools impacts peptidoglycan integrity. Since acetate is a weak inhibitor of Ddl, mechanisms that maintain a high intracellular D-Ala pools are sufficient to counter the effect of acetate-mediated Ddl inhibition in *S. aureus*.

---

Organic acids produced by host and bacterial metabolism are critical determinants of infection outcomes (1, 2). During infection, the host macrophages produce millimolar amounts of itaconate, a dicarboxylic acid known to inhibit bacterial growth (3). Conversely, many bacterial pathogens and the gut microflora excrete short-chain organic fatty acids, which exhibit immunomodulatory functions and can skew the host response during infection (4, 5). Upon entry into the bacterial cell, organic acids can become toxic to bacteria when they dissociate in the cytosol as protons and organic anions. The influx of protons can result in cytoplasmic acidification and prove lethal for some pathogens if not adequately controlled (6). Similarly, organic anions have been shown to accumulate to toxic levels in the bacterial cytoplasm (7). However, the precise consequences of organic anion toxicity and the mechanisms pathogens employ to withstand the effects of anion perturbations within cells are not clearly understood.

Here, we focus on the response of *Staphylococcus aureus* to acetic acid, which is the primary end-product of glucose catabolism under aerobic conditions. *S. aureus* also likely encounters high concentrations (up to 100 mM) of acetic acid and other short-chain fatty acids produced by human gut microbiota during intestinal colonization (8–10). On average, 20% of adults carry *S. aureus* in their intestines (11), and the burden there often surpasses that found in nasal passages by more than three orders of magnitude, establishing the gut as a primary site for *S. aureus* colonization (12). We have previously shown that excreted acetic acid can promote cytoplasmic acidification in cultures of *S. aureus,* especially when the external environment becomes sufficiently acidic (pH< 5) (13). Cytoplasmic acidification promotes protein oxidation and triggers a staphylococcal ClpP-dependent damage response that eliminates unfit cells from the population (14). In contrast, in mildly acidic environments (pH 5.5-6.5), although *S. aureus* actively buffers its intracellular environment against acidification, the transmembrane pH gradient (ΔpH) of *S. aureus* will drive the accumulation of millimolar quantities of acetate anions into the cytoplasm. Previous studies in *Escherichia coli* have shown that acetate intoxication causes an osmotic imbalance that can transiently be accommodated by the efflux of physiological anions like glutamate (7). In addition, acetate anions have also been reported to impact enzymes in the methionine biosynthetic pathway, resulting in a toxic accumulation of homocysteine and a reduction in intracellular methionine leading to growth inhibition of *E. coli* (15, 16). However, it remains unclear if these effects are common to other bacteria.

Here, we demonstrate that the primary target of acetate intoxication in *S. aureus* is Ddl. Ddl is essential for staphylococcal growth and produces D-Ala-D-Ala dipeptide which is incorporated into peptidoglycan to cross-link the peptide side chains of neighboring glycan strands. We also demonstrate that carbon flux through alanine racemase and a tight control of Dat activity increases the cytosolic D-Ala pool to counter acetate-mediated inhibition of Ddl. Importantly, these phenotypes are not unique to acetate, but are conserved across multiple biologically important organic acid anions. Therefore, we propose that *S. aureus* may have evolved to maintain a high intracellular D-Ala pool, partly to offset the inhibition of Ddl by organic anions typically encountered during human colonization.

## Results

### Alanine racemase counters acetate intoxication

To identify genetic determinants that counter the effects of acetate intoxication, we screened the Nebraska Transposon Mutant Library (NTML) for mutants sensitive to 20 mM acetic acid in Tryptic Soy Broth (TSB) media, pH 6.0. Under these conditions, *S. aureus* maintains its intracellular pH approx. 1.5 units above the external pH (17) and is estimated to accumulate over 600 mM acetate in the cytosol (18). The NTML strains were grown under static conditions at 37°C, and the extent of growth was determined at 24 h by measuring the optical density at 600 nm (OD_600_). As a control, we performed an identical screen without acetic acid supplementation. We normalized the growth of each mutant in both screens (± acetic acid) to their isogenic wild-type (WT) strain. A comparison of growth indices (OD_600_ Tn-mut/WT) for each mutant in the presence and absence of 20 mM acetic acid revealed that most mutants clustered close to an index of 1 in the plot (Figure 1A), which suggested that most mutants tolerated acetate intoxication reasonably well. A few mutants that grew poorly following acetate intoxication due to inherent growth defects were observed close to the plot diagonal, whereas those mutants that did not have intrinsic growth deficiencies were located further away from the diagonal. Among the latter class of mutants, we observed that the *alr1* mutant (SAUSA300_2027) had the most substantial reduction in growth when subjected to acetate stress (Figure 1A, B), with the severity of the phenotype depending on the acetate concentration (Figure 1C). To confirm that the acetate-dependent growth defect of the *alr1* mutant was not due to polar effects, we complemented the mutant by inserting a functional copy of *alr1* under the control of its native promoter into the *S. aureus* pathogenicity island (SaPI) attachment site. Genetic complementation completely restored the *alr1* mutant phenotype to WT levels (Figure 1B, C). These results suggest that acetate intoxication impairs the growth of *S. aureus* in the absence of a functional alanine racemase. Further supporting this conclusion, we could reduce acetate toxicity in the *alr1* mutant by culturing this strain in glucose-free TSB media, which alleviates carbon catabolite repression and activates TCA cycle-dependent acetate catabolism (Figure 1D) (19). Conversely, the inactivation of citrate synthase (*citZ*), the first enzyme of the TCA-cycle, re-imposed acetate toxicity in the *alr1* mutant when cultured in glucose-free TSB media (Figure 1D).

**Figure 1.**
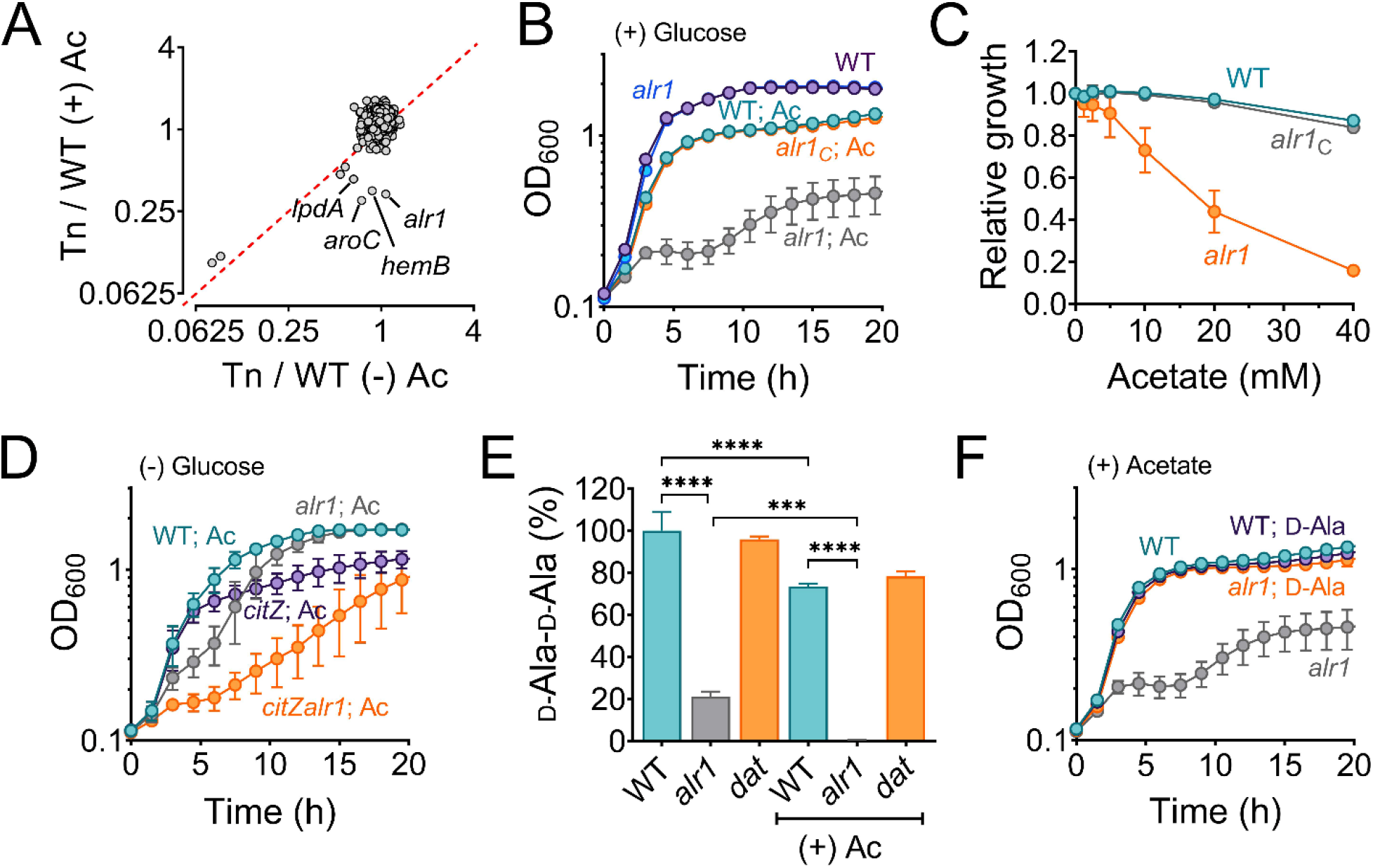
Alanine racemase activity counters acetate intoxication. **(A)** The Nebraska Transposon Mutant library was screened against 20 mM acetic acid, pH 6.0 to identify mutants with altered growth phenotypes. The WT strain and transposon mutants were grown for 24 h in TSB ± 20 mM acetic acid. The bacterial growth at 24 h was measured spectrophotometrically (OD_600_) and normalized to WT growth. The X and Y-axis on the plot represent normalized growth values for each mutant in the presence or absence of acetate. **(B)** The growth of the WT, *alr1* mutant, and *alr1* complemented strain (*alr1C*) in TSB supplemented with 20 mM acetic acid. **(C)** The tolerance of strains to various acetate concentrations was assessed by monitoring growth (OD_600_) following a previously published method (58). To maintain a transmembrane pH gradient of ∼1.5, the culture media was adjusted to pH 6, prior to challenging with subinhibitory acetate concentrations (40 mM-1.25 mM; two-fold dilutions). The relative growth (fractional area) of both the WT, *alr1* and *alr1C* mutant was calculated by comparing the area under the growth curves at subinhibitory concentrations of acetate to their corresponding controls (no acetate) and plotted against acetate concentrations. **(D)** Aerobic growth of WT, *alr1*, *citZ*, *citZalr1* mutants in TSB media lacking glucose, but supplemented with 20 mM acetic acid. **(E)** LC-MS/MS analysis was performed to quantify the intracellular D-Ala-D-Ala pool in strains cultured for 3 h (exponential phase) in TSB ± 20 mM acetic acid. **(F)** The growth of strains was monitored following D-Ala supplementation (5 mM) in TSB + 20 mM acetate, (n=3, mean ± SD). Ac, acetate. ***, P value <0.001; ****, P value <0.0001.

### Acetate intoxication alters the intracellular D-Ala-D-Ala pools

Alr1 catalyzes the conversion of L-Ala to D-Ala during staphylococcal growth (Figure 1-figure supplement 1A). The D-Ala is further converted to D-Ala-D-Ala dipeptide by the ATP-dependent Ddl (Figure 1-figure supplement 1A) and incorporated into peptidoglycan (PG) muropeptide, thus playing a crucial role in PG biosynthesis, cross-linking, and integrity (20, 21). Therefore, we hypothesized that under acetate stress, low concentrations of D-Ala in the *alr1* mutant might concomitantly reduce D-Ala-D-Ala concentrations in the cell resulting in a growth defect. To test this hypothesis, we determined the intracellular pool of D-Ala-D-Ala using liquid chromatography- tandem mass spectrometry (LC-MS/MS). In regular growth media (TSB), we observed that the inactivation of *alr1* decreased the D-Ala-D-Ala pool by approximately 80% compared to the WT strain (Figure 1E). However, following acetate intoxication, the level of D-Ala-D-Ala was depleted by more than 99% (Figure 1E). The external supplementation of D-Ala (5 mM) in the media fully restored the growth of the *alr1* mutant to WT levels under acetic acid stress (Figure 1F), which suggests that increased intracellular D-Ala pools can overcome the detrimental impact of acetate intoxication.

The depletion of D-Ala-D-Ala following acetate intoxication is surprising since *S. aureus* is predicted to have two additional pathways that can synthesize D-Ala and channel it to the production of this dipeptide. For instance, *S. aureus* harbors a second predicted alanine racemase (Alr2) that could compensate for the lack of Alr1 activity (Figure 1-figure supplement 1A). Alternatively, Dat, which catalyzes the formation of D-Ala from pyruvate and D-glutamate (D- Glu), may functionally complement the *alr1* mutant under acetate stress (Figure 1-figure supplement 1A). However, the lack of functional complementation from these alternate pathways of D-Ala biosynthesis following acetate intoxication suggests that not all metabolic routes to D- Ala are operational, or that regulatory bottlenecks limit pathway activity. To test these possibilities, we constructed a series of mutants in which all three predicted routes of D-Ala biosynthesis (*alr1*, *alr2* and *dat*) were disrupted either individually or in various combinations and performed growth assays (Figure 1-figure supplement 1B). Surprisingly, we observed that the inactivation of *alr1* and *dat* simultaneously (*alr1dat* mutant) was synthetic lethal in *S. aureus*, suggesting that *alr1* and *dat* were the sole contributors of D-Ala in *S. aureus*. Indeed, the supplementation of D-Ala fully restored the growth of the *alr1dat* mutant (Figure 1-figure supplement 1C).

The inactivation of *alr2,* either alone or in combination with other D-alanine-generating enzymes, did not affect growth (Figure 1-figure supplement 1B). This observation suggests that *alr2* is unlikely to be a functional alanine racemase under the growth conditions tested. Collectively, these results indicate that Dat activity accounts for D-Ala production in the absence of *alr1*, but its contribution is insufficient to counter acetate intoxication.

### Insufficient translation of *dat* impacts the *alr1* mutant following acetate intoxication

Since Dat activity contributes to D-Ala production in the *alr1* mutant, we questioned why Dat is insufficient to sustain D-Ala-D-Ala pools under conditions of acetate intoxication. One possible explanation may relate to the maintenance of osmotic balance by *S. aureus*. It has been proposed that the intracellular accumulation of acetate anions may bring about an efflux of L/D-Glu from cells to adjust for osmolarity, thus exhausting one of the key substrates for Dat activity and limiting D-Ala production (7). However, this hypothesis is improbable since the expression of *dat* from a multicopy vector rescued the *alr1* mutant from the effects of acetate intoxication (Figure 2-figure supplement 1A), suggesting that the intracellular D-Glu pools are sufficient to support D-Ala production through Dat activity. Alternatively, we hypothesized that the *alr1* mutant’s heightened sensitivity to acetate toxicity could be due to a decrease in *dat* transcription which would effectively reduce intracellular D-Ala. However, we found no detrimental effect of acetate intoxication on *dat* transcription in the *alr1* mutant (Figure 2-figure supplement 1B). Together, these observations raise the possibility that the depletion of D-Ala-D-Ala in the *alr1* mutant following acetate intoxication may arise from a post-transcriptional regulatory bottleneck, that limits *dat* from meeting the demand for intracellular D-Ala.

In *S. aureus*, *dat* is part of a bicistronic operon (Figure 2A). The first gene, *pepV*, encodes an extracellular dipeptidase (22, 23). Transcriptional start site (TSS) mapping of the *pepV-dat* operon by the adaptor and radioactivity-free (ARF-TSS) method revealed a 30-nucleotide untranslated region (5’-UTR) extending upstream from the *pepV* initiation codon. The 5’-UTR includes a Shine-Dalgarno motif (ribosome binding site, SD1) upstream of the *pepV* start codon (Figure 2A). In addition, a second SD motif (SD2) associated with *dat* was identified within the *pepV* coding region (Figure 2A), and did not overlap with the *pepV* termination codon. The location of SD2 within *pepV* suggests that the insufficient production of D-Ala by *dat* following acetate intoxication could be attributed to suboptimal translation of *dat*. This could occur as ribosomes (70S) that are moving from SD1 may interfere with the translation of *dat* from SD2. To test this hypothesis, we engineered a nonsense mutation in *pepV* (*alr1pepV^Q12STOP^* mutant) that would prevent the ribosomes originating from SD1 from moving forward (Figure 2A). However, the *alr1pepV^Q12STOP^* mutant grew poorly compared to the *alr1* mutant following acetate intoxication (Figure 2B). This suggested that the translation of *dat* is coupled to that of *pepV* presumably through stable mRNA secondary structures that form within *pepV*. These structures may not be effectively resolved in the *alr1pepV^Q12STOP^* mutant due to the absence of ribosome traffic on *pepV* mRNA.

**Figure 2.**
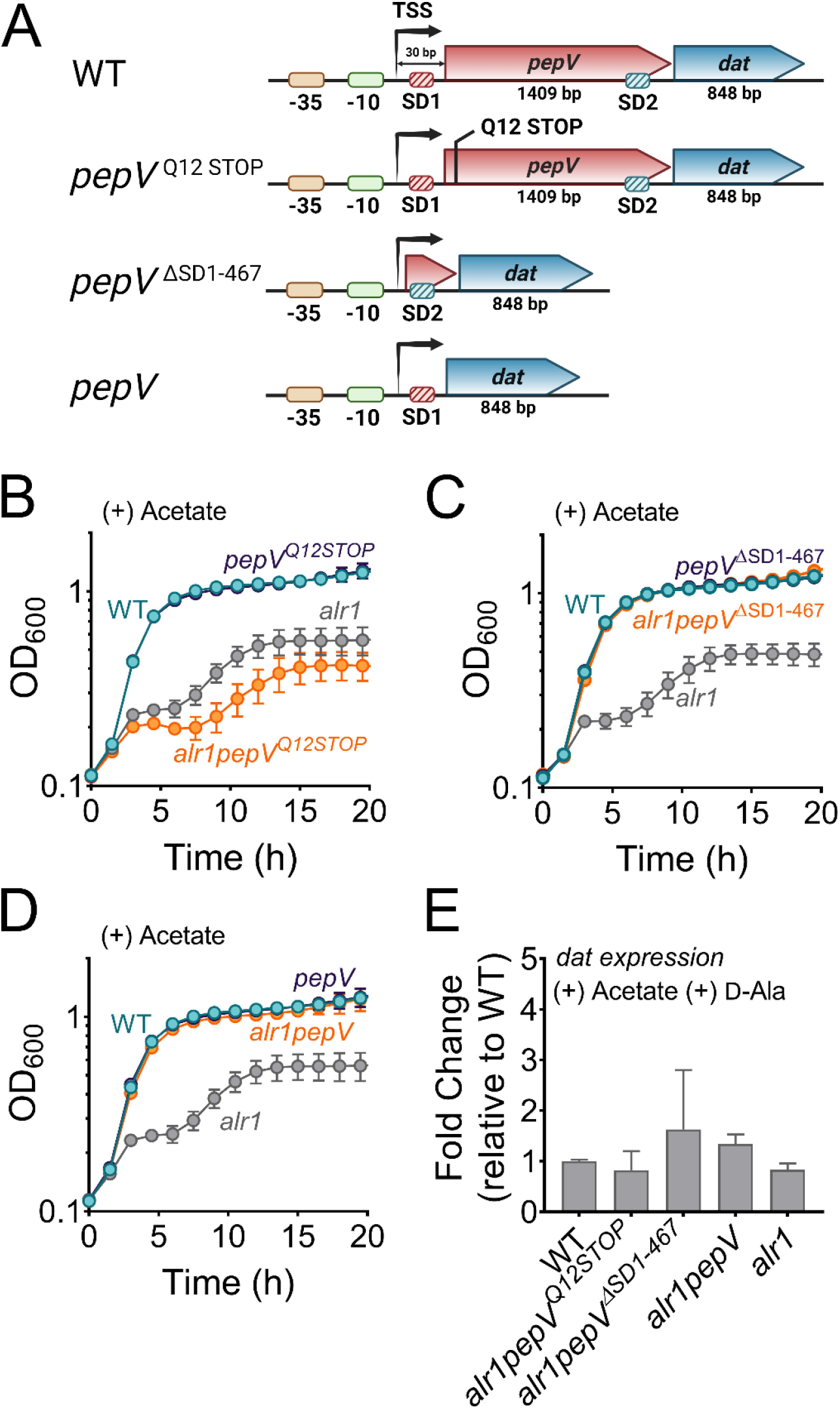
Translational coupling of *dat* to *pepV* limits the *alr1* mutant from countering acetate intoxication. **(A)** Schematic representation of various engineered mutations in the *pepV*- *dat* locus.SD, Shine-Dalgarno motif; TSS, transcriptional start site. **(B)-(D)** Growth of engineered mutants was monitored spectrophotometrically (OD_600_) in TSB supplemented with 20 mM acetate (n=3, mean ± SD). **(E)** RT-qPCR to determine *dat* transcription in various mutants relative to the WT strain.

As an alternative approach to determine if SD2 positioning within *pepV* impeded *dat* translation, we deleted *pepV* along with SD1 in the *alr1* mutant (*alr1pepV*^ΔSD1-467^, Figure 2A). In the resulting strain, *dat* translation was under the sole control of its native SD2. Remarkably, the *alr1pepV*^ΔSD1-467^ mutant did not display a heightened sensitivity to acetate stress and grew identical to the WT strain following acetate intoxication (Figure 2C). Similarly, an *alr1* mutant in which *dat* was linked to SD1 (*alr1pepV* mutant, Figure 2A) also phenocopied the WT strain following acetate intoxication (Figure 2D). Notably, the observed growth differences in *alr1pepV*^ΔSD1-467^, *alr1pepV* and *alr1pepV^Q12STOP^*mutants following acetate intoxication did not result from any changes in *dat* transcription (Figure 2E). These findings collectively suggest that the native promoter elements, as well as the SD sites of *pepV* and *dat* can independently support the robust expression and translation of *dat* to levels required for countering acetate intoxication. However, the genetic arrangement of the *dat* translation initiation region (TIR) within *pepV*, offered tight control of *dat* translation, and prevented cells from producing sufficient enzyme following acetate intoxication.

### Why is the Dat tightly controlled?

The need to tightly control Dat activity suggests that flux between D-Ala and D-Glu pools must be carefully balanced during staphylococcal growth. To gain insight into this process, we profiled the mass isotopologue distribution (MID) of D-Ala-D-Ala in the WT, *alr1,* and *dat* mutants under isotopic steady-state conditions using ^13^C ^15^N -L-Ala as the tracer during growth experiments in chemically defined medium (CDM). The flux of ^13^C3^15^N1-L-Ala through Alr1 should result in ^13^C ^15^N -D-Ala production (Figure 3A, D-Ala retains labeled nitrogen). On the other hand, staphylococcal alanine dehydrogenases (Ald1 and Ald2) catalyze the conversion of ^13^C ^15^N -L- Ala to ^13^C3-pyruvate, and finally ^13^C3-D-Ala through Dat activity (Figure 3A). Thus, the labeled nitrogen in ^13^C ^15^N -L-Ala is lost as ^15^N -NH when fluxed through the Ald/Dat pathway (Figure 3A). Since the intracellular pools of D-Ala are converted to D-Ala-D-Ala, the MID of the latter metabolite should mirror the isotopologue ratios of D-Ala produced from either Alr1 or Dat activities.

**Figure 3.**
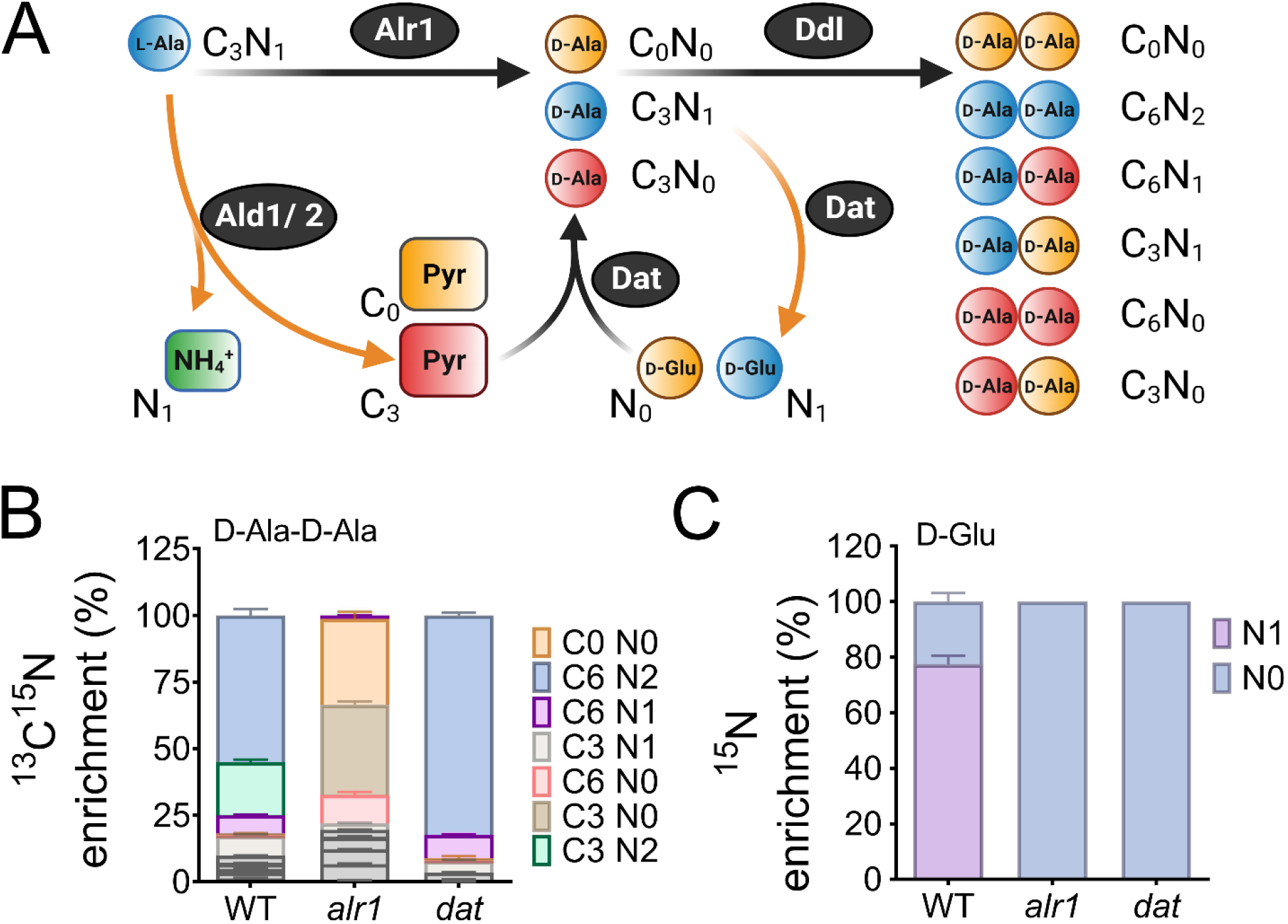
Reaction orientation and fluxes through Alr1 and Dat. **(A)** Schematic representation of various isotopologues of D-Ala-D-Ala and D-Glu generated from ^13^C3^15^N1 labeled L-Ala. Metabolites in blue mainly arise from Alr1, red, through the Ald1/2-Dat pathway and yellow are unlabeled intermediates within cells. The mass isotopologue distribution of **(B)** D-Ala-D-Ala and **(C)** D-Glu were determined by LC-MS/MS following the growth of *S. aureus* in chemically defined media supplemented with ^13^C3^15^N1 L-Ala (n=3, mean ± SD). Isotopologues of D-Ala-D-Ala shown in grey color are minor species and are noted in Table S9.

LC-MS/MS analysis revealed that ∼80% of the intracellular D-Ala-D-Ala pool had incorporated the labeled L-Ala supplemented in media (fractional contribution, 0.80). As expected, the majority (∼55%) of the D-Ala-D-Ala in the WT was composed of the C6N2 isotopologue (in which both units of D-Ala contain labeled carbon and nitrogen), which suggested that *alr1* was the major contributor of D-Ala in *S. aureus* (Figure 3B). Surprisingly, the sole contribution of *dat* activity (C6N0, C3N0, C0N0) to D-Ala-D-Ala was less than 1% in the WT strain, and D-Ala-D-Ala isotopologues with at least one D-Ala originating from *dat* activity (C6N1, C3N1, C3N2) although readily observed, were still in the minority. However, the D-Ala-D-Ala originating from Dat activity expanded substantially upon *alr1* mutation (Figure 3B). Inactivation of *dat* itself displayed few differences in the MID of D-Ala-D-Ala, compared to the WT strain (Figure 3B). These results suggest that flux through Dat is most likely driven towards D-Glu in the WT strain rather than D-Ala. Only upon inactivation of *alr1* does the Dat activity reverse towards the production of D-Ala. To confirm these predictions, we measured the levels of ^15^N1-D-Glu in the WT, *alr1*, and *dat* mutants following growth with the ^13^C3^15^N1-L-Ala tracer. Consistent with Dat activity funneling D-Ala to D-Glu in the WT, approximately 78% of the D-Glu pool in the WT strain was ^15^N labeled. Furthermore, we observed that inactivation of *dat* resulted in the complete depletion of intracellular levels of ^15^N1-D-Glu (Figure 3C). Inactivation of *alr1* also had a similar outcome with loss of ^15^N1-D-Glu pools due to the lack of ^13^C ^15^N1-D-Ala in this mutant (Figure 3C). Together, these results strongly suggest that in the WT strain, Dat activity diverts D-Ala towards D-Glu production.

Given the critical need to produce D-Ala-D-Ala during acetate intoxication, any diversion of its precursor pool (D-Ala) to D-Glu through Dat activity is bound to decrease cell fitness and thus may justify its tight translational control. To test this hypothesis, we determined the mean competitive fitness (*w*) of cells that overexpressed *dat* compared to those that had native levels of expression. Accordingly, we performed coculture competition assays of the WT strain with an isogenic mutant strain that either harbored an empty vector (pAQ59) integrated into the SaPI chromosomal site or a vector containing *dat* under control of its native promoter (pAS8), following acetate intoxication. Consistent with increased Dat activity in the WT strain being detrimental to the cell, the mean competitive fitness of the *dat* overexpressing strain was significantly lower (*w*4h= 0.91) in the exponential growth phase than its isogenic WT strain that harbored the empty vector (*w*4h= 1.26) (Figure 3D). Collectively, these results suggest that Dat catalyzes the production of D-Glu in the WT strain, and its tight regulation prevents excessive flux of D-Ala to D-Glu which is necessary to maintain cell fitness following acetate intoxication.

### Acetate intoxication impacts PG biosynthesis

Since acetate intoxication ultimately affects D-Ala-D-Ala pools (Figure 1D), we predicted potential alterations to PG biosynthesis and cell wall integrity. To test this hypothesis, we quantified various cytosolic PG intermediates in the WT strain by LC-MS/MS analysis. Acetate intoxication caused a significant increase in the intracellular pools of multiple PG biosynthetic intermediates, including Uridine diphosphate N-acetylglucosamine (UDP-NAG), UDP-N-acetylmuramic acid (UDP-NAM), UDP-NAM-L-Ala, UDP-NAM-L-Ala-D-Glu-L-Lys and UDP-NAM-L-Ala-D-Glu-L-Lys-D-Ala-D-Ala (UDP-NAM-AEKAA) in the WT strain when compared to the unchallenged control (Figure 4A). However, the growth of the WT strain was slightly inhibited by acetic acid (Figure 1B), which suggests that the observed accumulation of PG intermediates may have been due to an imbalance between the rates of PG biosynthesis and growth. Notably, the *alr1* mutant showed higher levels of UDP-NAM-AEK compared to the WT and the *dat* mutant following acetate intoxication (Figure 4A), indicating a metabolic block in the production of UDP-NAM-AEKAA due to insufficient D-Ala-D-Ala. The effect of this metabolic block is also evident from the increased transcription of *ddl* and *murF* (Figure 4B) which encode enzymes that incorporate D-Ala-D-Ala into PG precursors, suggesting a greater need to maintain peptidoglycan cross-linking following acetate intoxication.

**Figure 4.**
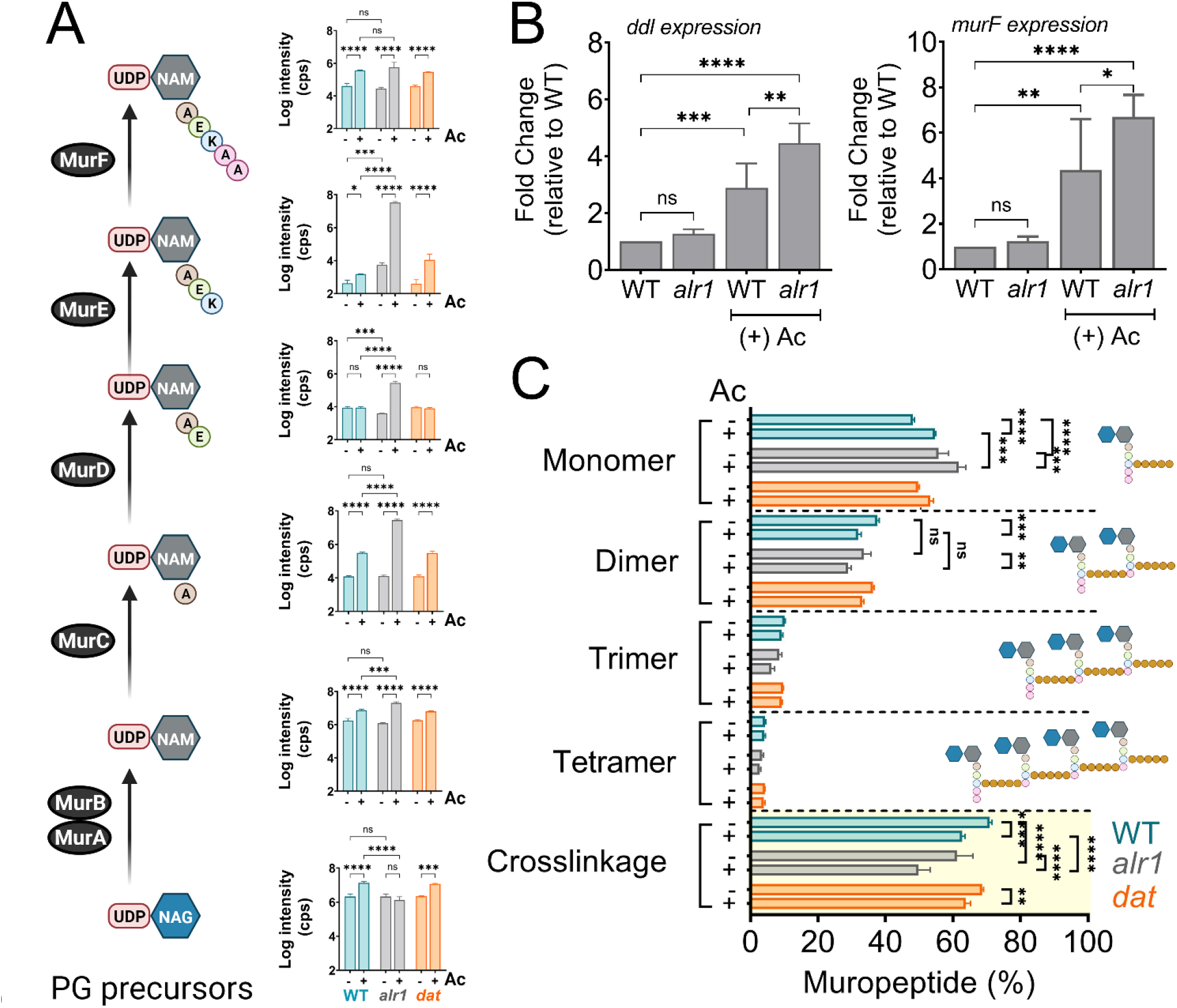
Acetate intoxication impacts soluble PG precursor pools and cell wall cross- linking. **(A)** The intracellular pool of PG intermediates in exponential phase cultures of *S. aureus* was estimated using LC-MS/MS analysis. cps, counts per second **(B)** *ddl* and *murF* transcription in the exponential growth phase was determined by RT-qPCR analysis (n=3, mean ± SD). **(C)** Cell wall muropeptide analysis of the WT, *alr1* and *dat* mutants was determined following growth in TSB ± 20 mM acetate for 3 h. Cell wall cross-linking was estimated as previously described (53). Ac, acetate. *, P value <0.05; **, P value <0.01; ***, P value <0.001; ****, P value <0.0001.

Unsurprisingly, the dysregulation of D-Ala-D-Ala homeostasis following acetate intoxication was also reflected in the extent of cell wall cross-linking in the WT, *alr1* and *dat* mutants. Muropeptide analysis revealed that acetate intoxication in the WT strain increased levels of monomeric muropeptides (Figure 4C, Figure 4-figure supplement 1). Conversely, the percentage of di- and trimeric muropeptides decreased relative to the WT control, as did the percent cross- linking (Figure 4C). These observations suggest that acetate intoxication constrains the D-Ala-D-Ala pool in the WT strain and alters PG cross-linking despite Alr1 activity. The extent of PG cross- linking in the *dat* mutant was similar to WT in the presence or absence of acetate, consistent with our finding that the Dat activity plays a limited role in maintaining the D-Ala-D-Ala pool in the WT strain (Figure 4C). In contrast, PG cross-linking in the *alr1* mutant was lower than the WT strain by ∼10% (Figure 4C). Acetate intoxication further decreased the cross-linking approximately 20% relative to WT as well as the ratio of dimeric to monomeric muropeptides in the *alr1* mutant, which inevitably reduced the growth of this strain (Figure 4C).

Muropeptide analysis also revealed the accumulation of a disaccharide tripeptide (NAG-NAM-AEK (M3); m/z. Da, 826.4080) in the peptidoglycan (PG) extracted from the *alr1* mutant (Figure 4-figure supplement 1B). This finding suggests that the significantly elevated levels of UDP-NAM-AEK in the *alr1* mutant could efficiently outcompete the substrate specificity of phospho-N-acetylmuramyl pentapeptide translocase (MraY) for UDP-NAM-AEKAA, ultimately becoming integrated into the PG structure itself. Interestingly the incorporation of UDP-NAG-NAM-AEK into the *alr1* mutant’s PG only marginally increased following acetate treatment (Figure 4-figure supplement 1B, see inset). The increase of UDP-NAG-NAM-AEK is most likely an underestimate since cells with higher levels of incorporation are more likely to lyse due to a reduction in PG cross-linking. Overall, these observations support a model wherein the immediate consequences of acetate intoxication are defects in PG crosslinking and biosynthesis.

### Acetate intoxication inhibits Ddl activity

While the above observations point to the consequences of acetate intoxication of *S. aureus*, its molecular target was not initially identified. Since acetate intoxication dramatically reduces D-Ala-D-Ala levels in the *alr1* mutant (Figure 1D), we reasoned that acetate might inhibit either Dat or Ddl activity. To distinguish between these two targets, we measured the levels of D-Ala in the *alr1* mutant following acetate intoxication. Surprisingly, we observed that the D-Ala pools in the *alr1* mutant did not significantly change in response to acetate intoxication compared to the untreated control (Figure 5A). This suggested that Dat activity was preserved in the *alr1* mutant to the same extent as its untreated control and was not affected by acetate.

**Figure 5.**
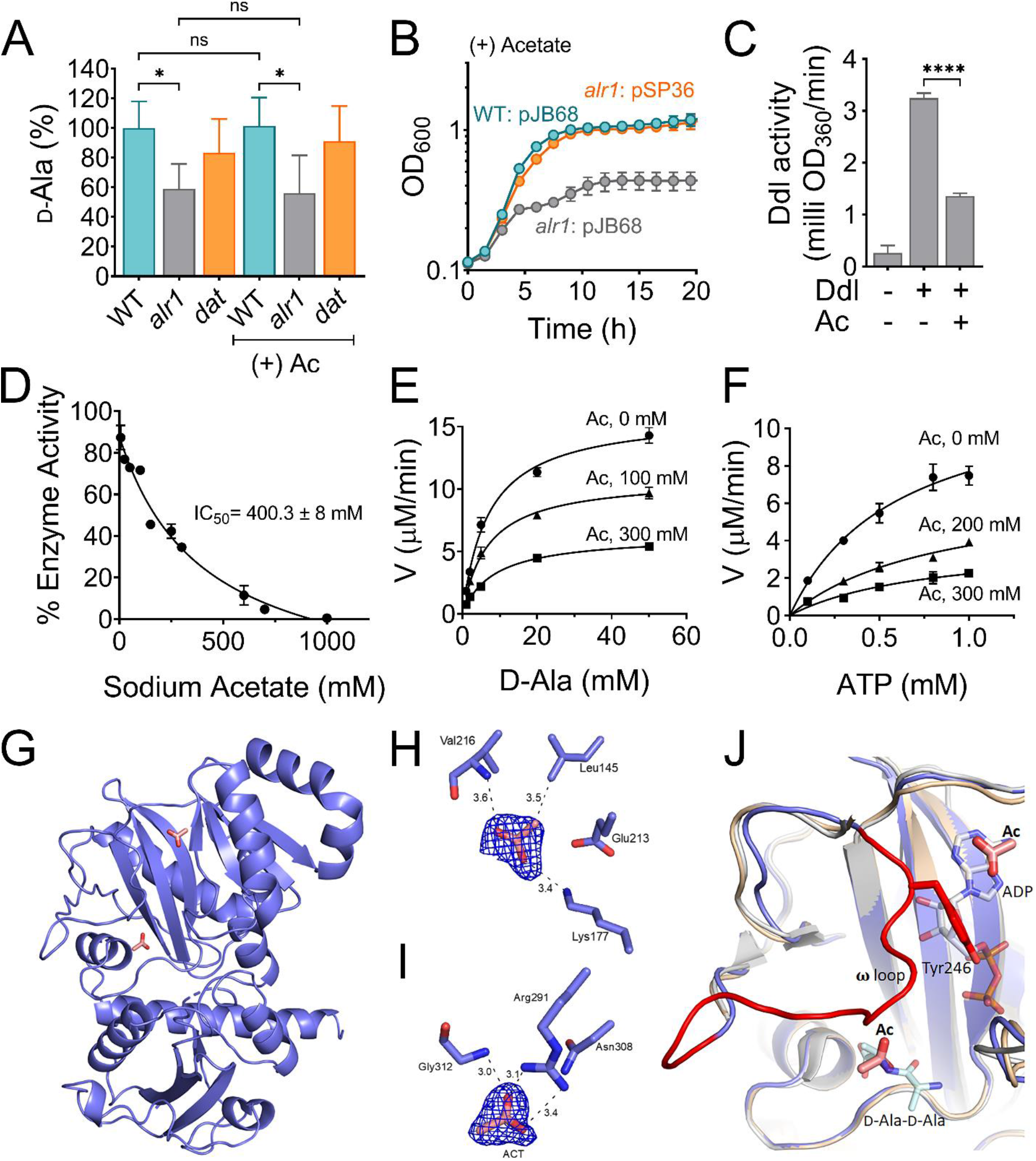
Acetate anion inhibits Ddl activity. **(A)** The intracellular D-Ala was determined by LC- MS/MS analysis. **(B)** The *ddl* gene was overexpressed in *S. aureus* using a cadmium inducible expression system (pSP36). CdCl2, 0.312 µM. **(C)** Inhibition of recombinant His-tagged Ddl activity in the presence of 300 mM sodium acetate **(D)** IC50 curve of the inhibition of rDdl by acetate. Michaelis-Menten kinetics of rDdl in varying concentrations of **(E)** D-Ala, and **(F)** ATP in the presence of acetate to assess the inhibition mechanism. **(G)** Structure of the acetate bound Ddl (PDB:8FFF). **(H)** Acetate bound to the ATP binding site of Ddl **(I)** Acetate bound to the second D-Ala binding site of Ddl. The calculated Fo-Fc omit maps are contoured to 3σ and the mesh is shown in blue. **(J)** Superimposed structure of acetate bound Ddl (slate blue) with StaDdl apo structure (PDB:2I87, beige) and StaDdl-ADP complex structure (PDB:2I8C, grey) showing a shift of ω loop (red) to ATP binding site. The D-Ala-D-Ala was modeled at the D-Ala binding site using *Thermos thermophius* HB8 Ddl structure (PDB:2ZDQ). The bound ADP (grey) of PDB:2I87 and modeled D-Ala-D-Ala (light blue) indicates the positioning of Ac at ATP and second D-Ala binding sites respectively. Ac, acetate; V, velocity; *, P value <0.05; ****, P value <0.0001.

Conversely, these findings also indicate that the acetate-dependent decrease of the D-Ala-D-Ala pool in the *alr1* mutant was most likely due to the inhibition of Ddl. To test this hypothesis, we cloned *S. aureus ddl* under the control of a cadmium inducible promoter and induced its expression in the *alr1* mutant following acetate intoxication (Figure 5B). Indeed, the growth of the *alr1* mutant was restored to WT levels when *ddl* was overexpressed, strongly suggesting that Ddl was the target of acetate anion (Figure 5B).

To confirm that acetate inhibits Ddl through direct interactions, we undertook two separate approaches. As the first approach, 6xHis-tagged *S. aureus* Ddl was purified, and in-vitro enzyme kinetic assays were performed to determine the possible inhibitory mechanism of Ddl by acetate.

Considering the high concentration of acetate estimated to accumulate in the cytoplasm, a concentration of 300 mM sodium acetate was used in the initial reactions to test inhibition (Figure 5C). Interestingly, variation of acetate concentration showed that Ddl was inhibited *in vitro*, and these conditions suggest an IC50 of 400.3 ± 8 mM (Figure 5D). This indicates significant inhibition of Ddl by acetate when the cellular concentration is near the hypothesized 600 mM, further confirming that Ddl is a direct target of inhibition by acetate anion. Furthermore, based on kinetic experiments performed under varying concentrations of either ATP or D-Ala, the *k*cat values are shown to be distinctly different for each acetate concentration, which strongly suggests a mixed inhibition mechanism for acetate (Figure 5E and F, Table S1).

Differential Scanning Fluorometry (DSF) was used as another approach to assess the direct binding of acetate to Ddl (Table S2). Ddl has two D-Ala binding sites and one ATP-binding site (24), and DSF experiments were conducted with various combinations of these ligands. The Ddl protein without any ligand bound shows a melting temperature (Tm) of 45 °C. After adding 300 mM sodium acetate, Ddl exhibited a 3.7 °C Tm shift indicating a slight thermal stabilization upon binding acetate. This is higher than the shift in the Tm exhibited by a Ddl/ATP complex. The addition of ADP to Ddl results in a decrease of 3.2 °C, indicating a decrease in thermal stability compared to Ddl alone. Intriguingly, when adding acetate to Ddl complexes with ATP or ADP, the Tm increased to 48.9 °C and 49.9 °C, respectively (Table S2). This represents a Tm increase of 2.3 °C when acetate is added to a Ddl/ATP complex, but a Tm increase of 8.1 °C when acetate is added to a Ddl/ADP complex. The addition of D-Ala to the reaction mixture increases the Tm of Ddl by 4.2 °C and adding acetate to the Ddl/D-Ala mixture shows only a 0.3 °C Tm shift (Table S2). The widely varying changes in Tm for the tested complexes, particularly when comparing the Tm values for ligand-free, ADP-bound, and the ADP/Acetate complex, further support a mixed inhibition mechanism as these data suggest acetate may bind to multiple sites on Ddl or the location of these binding sites may change depending on the ligand-bound state of the enzyme due to Ddl conformational changes as observed in Ddl orthologs (24).

### Ddl/Acetate complex structure shows binding of acetate at both substrate binding sites

To gain further insight into the mechanism of acetate inhibition, the X-ray crystal structure of a Ddl/acetate complex was obtained using co-crystals of Ddl and acetate. The crystal diffracted to 1.9 Å and data were consistent with a P 2 21 21 space group possessing one molecule of Ddl in the asymmetric unit (Table S3). The crystal structure of the Ddl/acetate complex (PDB:8FFF) shows difference density corresponding to acetate at two different sites of the protein. One acetate is positioned within the adenine binding subsite of the ATP binding site and the other acetate ion is positioned in the second D-Ala binding site (Figure 5G). The acetate ion in the ATP binding site interacts with the side chain of Lys177 and the backbone nitrogen of Val216. Also, the methyl group of acetate forms van der Waals interactions with the side chain of Leu145 (Figure 5H). The acetate ion that binds to the D-Ala binding site forms a bidentate polar interaction with the side chain of Arg291 and a hydrogen-bonded interaction with the backbone nitrogen of Gly312 (Figure 5I). These two residues are conserved in Ddl homologs and previous structural data clearly illustrate the crucial role these residues play in D-Ala binding (24).

The acetate-bound structure shows conformational differences compared to the previously published ligand-free and ADP-bound structures (24). The ω loop, which is associated with substrate binding, is disordered in both the *S. aureus* Ddl ligand-free (PDB:2I87) and the Ddl- ADP complex structures (PDB:2I8C) as well as other available Ddl crystal structures that lack bound substrates or ligands (PDB:3K3P, 5DMX and 6U1C) (25–27). Interestingly, this loop is well ordered in the acetate-bound structure described here (Figure 5J), which gives the first view of the *S. aureus* Ddl ω loop and the interactions it may form with substrates or inhibitors. The structural stabilization of the ω loop is consistent with the DSF results exhibiting an increase in the melting temperature upon binding acetate. The ω loop is shifted towards the ATP binding site and repositions the conserved Tyr246 side chain within the ATP binding site, which likely hinders the binding of ATP (Figure 5J). This positioning is comparable with the *Mycobacterium tuberculosis* Ddl (PDB:3LWB) ligand-free structure, which also takes a closed conformation showing the ω loop positioned within the ATP binding site and obstructing ATP binding (28). Taken together, the kinetic, DSF, and structural data suggest that while acetate can directly bind within both substrate binding pockets of Ddl, it also stimulates conformational changes in the dynamic ω loop to afford more allosteric-like effects on enzyme activity. Each of these observations support a mixed inhibition modality.

### Multiple organic acids inhibit the *alr1* mutant in a D-Ala-dependent manner

Finally, we determined whether the growth inhibition of the *alr1* mutant is unique to acetate anion or is a more general phenomenon mirrored by addition of other small organic acids. Accordingly, we initially performed molecular docking studies of three biologically relevant organic anions: lactate, propionate and itaconate, in both the ATP and D-Ala binding pockets of Ddl (Figure 6A-D). The acetate anion-bound structure of Ddl was used as a reference for analysis. The docking results suggest reasonable poses for lactate, propionate and itaconate within the ATP binding site forming polar interactions with Ddl residues conserved for binding ATP. Upon docking, the carboxylate moieties of both lactate and propionate form ionic interactions with the Lys177 side chain similar to those observed in the Ddl/acetate crystal structure (Figure 6A and B). Also, the side chain of Glu213 in Ddl forms a hydrogen bonded interaction with the hydroxyl of lactate (Figure 6A) and van der Waals interactions between propionate and nearby side chains of Phe175 and Phe295 (Figure 6B) were indicated. The two carboxylate groups of itaconate form hydrogen bonded interactions with backbone amide nitrogen atoms of Ala218 and Tyr246 as well as a van der Waals interaction with the nearby side chain of Phe175 (Figure 6C).

**Figure 6.**
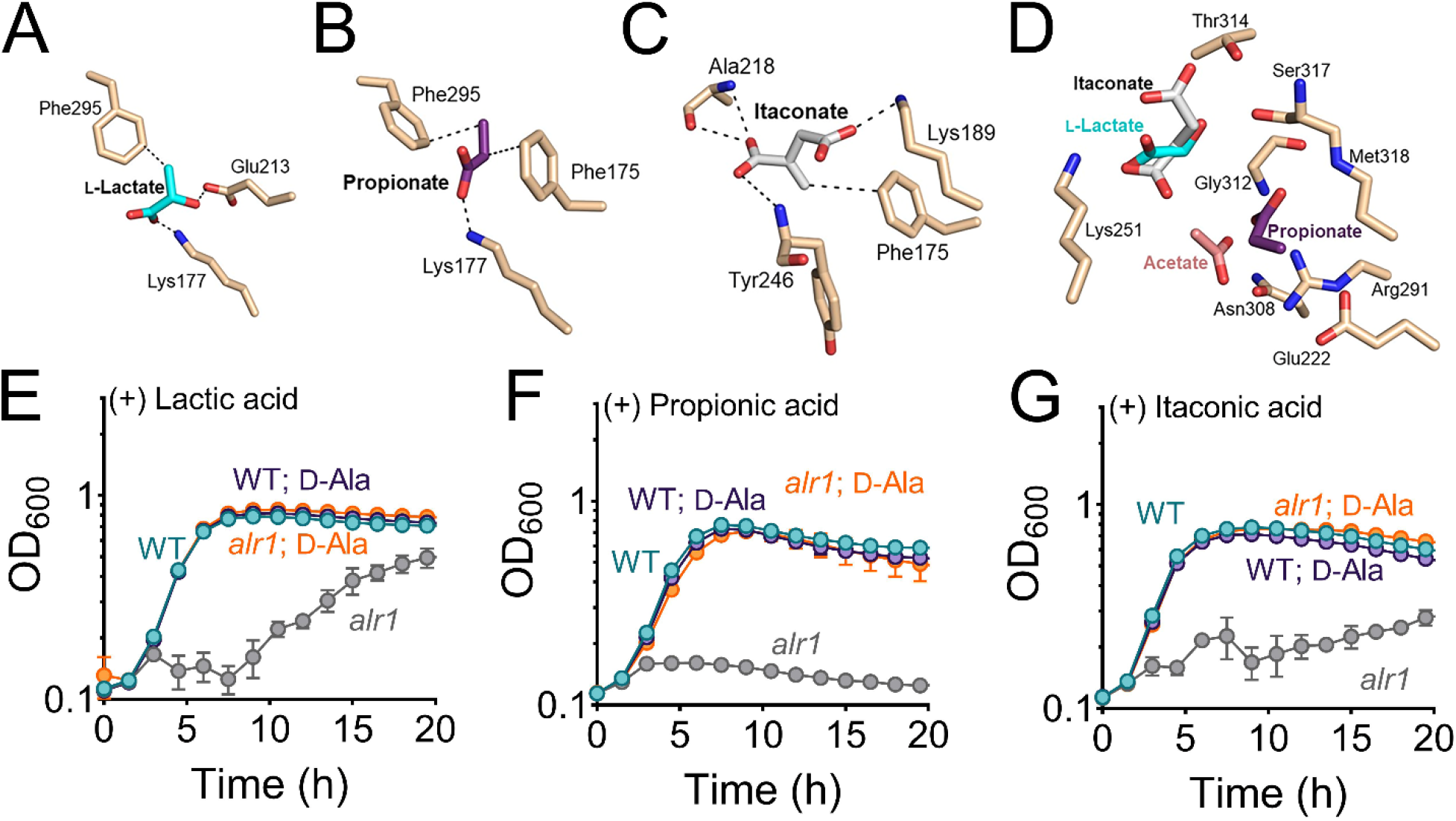
Biologically relevant weak acids inhibit growth of the *alr1* mutant. Molecular docking of **(A)** lactate **(B**) propionate and **(C)** itaconate to the ATP binding site of Ddl. **(D)** The relative positions and poise of different organic anions in relation to acetate in the D-Ala binding site of Ddl was determined using Schrödinger Glide. The growth (OD_600_) of the WT and *alr1* mutant in TSB containing **(E)** lactic acid (40 mM) **(F)** propionic acid (20 mM) and **(G)** itaconic acid (20 mM) in the presence or absence of 5 mM D-Ala.

The molecular docking results for lactate, propionate, and itaconate in the D-Ala binding site of Ddl also show similar types of interactions, but with variable poses and slight orientation differences compared to that observed for acetate in the crystal structure (Figure 6D). The D-Ala binding site, consisting of primarily charged and polar atoms, allows for a range of binding modes for these small anions, where the ligand size is a stronger factor in determining the binding location. Acetate and propionate, being smaller and less sterically hindered, bind preferentially near the Arg 291 side chain that coordinates the acid moiety of D-ala during the enzymatic reaction (Figure 6D). Meanwhile, itaconate and L-lactate bind in the more spacious region between Lys251 and Ser317 (Figure 6D). The Glide scores from the docking results, which provide a rough estimate of the ΔG of binding for each ligand suggest modest affinity to the identified binding sites (Table S4).

To determine if these organic acids could impact Ddl function, the WT and the *alr1* mutant were challenged with lactic, propionic and itaconic acids (Figure 6E-G). All three organic acids inhibited the growth of the *alr1* mutant. The addition of D-Ala to the culture media rescued the growth of the *alr1* mutant to WT levels, (Figure 6E-G) consistent with Ddl being the target of lactate, propionate and itaconate. Moreover, overexpression of *ddl* in the *alr1* mutant also restored growth of the *alr1* mutant following the organic acid challenge (Figure 6-figure supplement 1A-C). These findings collectively suggest that various organic acid anions can inhibit Ddl activity in *S. aureus*.

## Discussion

Intracellular anion accumulation has long been hypothesized to drive weak organic acid toxicity in bacteria (18, 29, 30). However, few studies have investigated the mechanism by which weak acid anions inhibit bacterial growth. Acetic acid is particularly interesting among weak acids, given that it is a common byproduct of glucose catabolism in bacteria and is excreted in high concentrations (31). *S. aureus* does not catabolize acetate as a carbon source unless glucose is first exhausted from its environment (32). Thus, in the presence of glucose, acetate can accumulate intracellularly in *S. aureus* as a function of the bacterial transmembrane pH gradient, especially when acetic acid concentrations are high in the immediate vicinity of cells. Here we determine that at high intracellular concentrations, acetate anions directly bind Ddl and inhibit D-Ala-D-Ala production to adversely impact peptidoglycan cross-linking (Figure 7).

**Figure 7.**
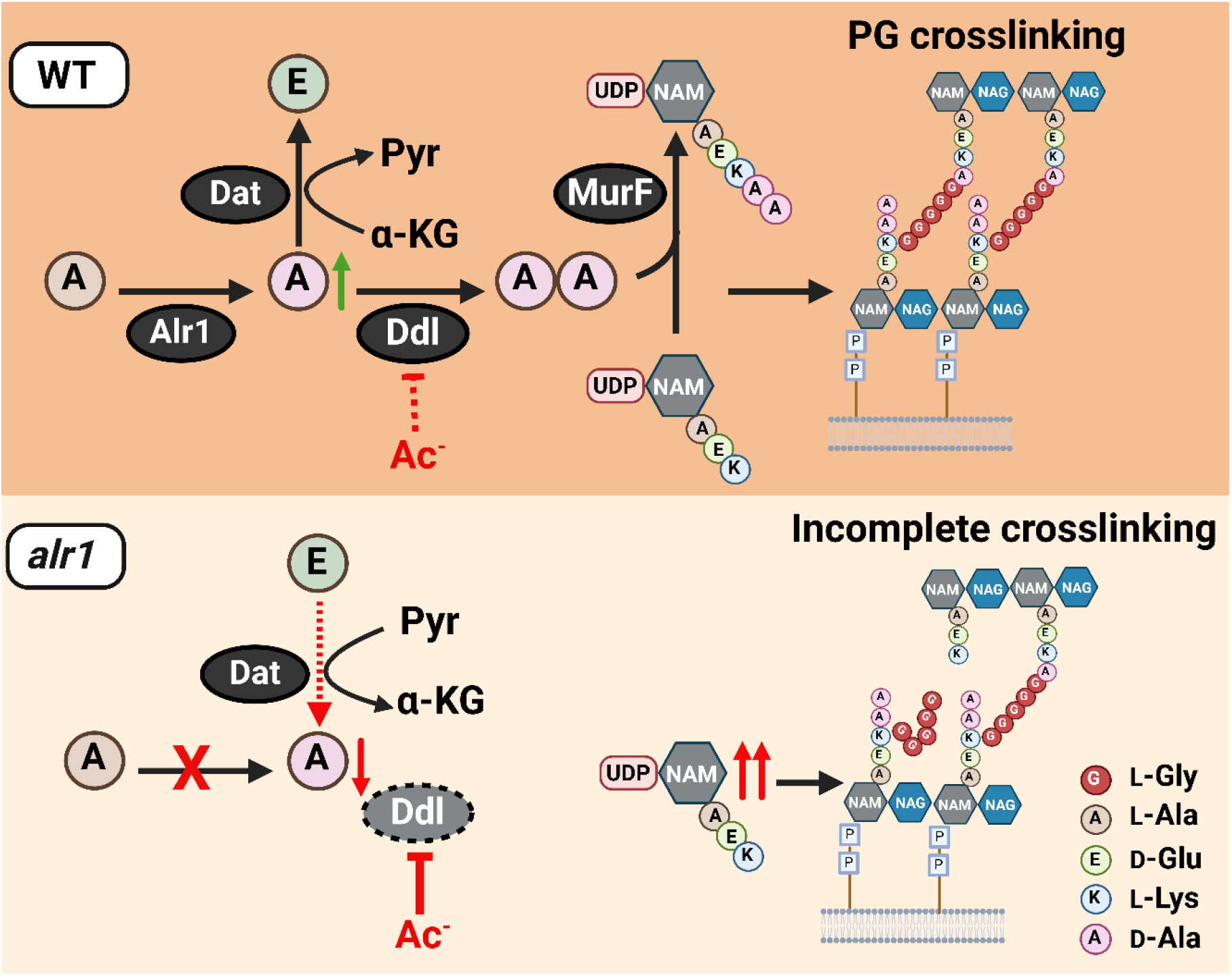
Model depicting the role of Alr1 in countering organic acid anion-mediated inhibition of Ddl. During its growth, *S. aureus* (WT) maintains a substantial intracellular pool of D-Ala through the activity of Alr1. Any excess D-Ala is subsequently converted into D-Glu by the action of the Dat enzyme. The high concentration of D-Ala is crucial for the optimal functioning of Ddl and serves to prevent the inhibition of Ddl by acetate (Ac^-^) and other organic acid anions. This process generates sufficient D-Ala-D-Ala, which is rapidly incorporated into the PG tripeptide precursor UDP-NAM-AEKAA to form UDP-NAM-AEKAA, which ultimately contributes to a robust cross-linked PG (murein) sacculus. In the *alr1* mutant, the Dat reaction orientation is switched to preserve intracellular D-Ala. Nevertheless, this change is inadequate to maintain sufficient D-Ala pool to shield Ddl from inhibition by Ac^-^, due to tight control of *dat* translation. This results in an excess of UDP-NAM-AEK, which competes effectively with UDP-NAM-AEKAA for PG incorporation. The absence of a terminal D-Ala-D-Ala in the PG hinders crosslinking and leads to impaired growth following acetate intoxication.

However, *S. aureus* exhibits a remarkable tolerance to acetate intoxication due to the robust production of D-Ala by Alr1, which ultimately increases D-Ala-D-Ala pools (Figure 7).

Multiple lines of evidence demonstrate Ddl to be the target of acetate anions. First, LC- MS/MS analysis revealed that acetate intoxication decreased D-Ala-D-Ala pools, but not D-Ala in *S. aureus*, pointing to Ddl as the target of acetate. Second, DSF and in-vitro enzyme kinetic studies showed that acetate could bind and inhibit purified rDdl through a mixed inhibition mechanism. Third, structural analysis of the Ddl-inhibitor complex confirmed that acetate binds to both the ATP-binding and D-Ala binding sites within Ddl and further induced conformational changes to the dynamic ω loop, which weakens the binding of ATP to the Ddl active site. Finally, overexpression of *ddl* alone was sufficient to overcome acetate-mediated inhibition of the *alr1* mutant and restore growth to WT levels.

Inhibitors that bind an enzyme’s catalytic substrate binding sites are usually competed out by high concentrations of substrates. However, acetate inhibits Ddl through a mixed inhibition mechanism despite binding to the substrate binding pockets of Ddl. We suspect this is due to additional conformational changes observed in the dynamic ω loop that affords more allosteric- like effects on enzyme activity. However, we cannot rule out that acetate might bind to additional sites in the Ddl-ATP complex, Ddl-ADP complex, or a Ddl-ADP-phospho-D-Ala complex with varying affinities. The differences in the temperature shifts observed in DSF with various substrate complexes support this possibility. The crystal structures of Ddl/acetate complexes with different substrates could provide a more precise conclusion about the inhibitory modality of Ddl by acetate. In line with acetate’s inhibitory effect on Ddl, we observed that acetate intoxication in the *alr1* mutant led to a disproportionate increase in the cytosolic pool of PG tripeptide intermediate (UDP-NAM-AEK) compared to the pentapeptide form (UDP-NAM-AEKAA). Previous reports have suggested that MraY might facilitate the integration of UDP-NAM- tripeptide into *S. aureus* PG, especially when its concentration within cells exceeds that of UDP-NAM-pentapeptide (33, 34). Our findings strongly support this hypothesis, as the analysis of the *alr1* mutant’s cell wall muropeptides revealed a clear elevation in the level of the disaccharide-tripeptide NAG-NAM-AEK. The inhibition of Ddl by acetate would further reduce the presence of terminal D-Ala-D-Ala moieties within *alr1* muropeptides which likely leaves these cells incapable of withstanding the outward-directed cell turgor pressure, ultimately leading to cell death (34).

Despite acetate inhibiting Ddl through a mixed inhibition mechanism, it should be noted that a functional Alr1 or even the supplementation of D-Ala in culture media can provide significant tolerance against acetate intoxication in *S. aureus*. These observations suggest that Ddl is only weakly inhibited by acetate, which is also evident from the relatively high IC50 of approximately 400 mM observed in our kinetic experiments with *S. aureus* Ddl. The weak inhibition of Ddl would suggest that inflating the cytosolic D-Ala pools could promote sufficient generation of D-Ala-D-Ala to counter acetate intoxication. Indeed, it has been estimated that *S. aureus* maintains a high concentration of roughly 30 mM intracellular D-Ala (35), which we now demonstrate to be critical in countering acetate intoxication. High concentrations of acetate and other short-chain fatty acids are typically found in the human gut, where *S. aureus* can colonize (8–10, 12). In these environments, the robust production of D-Ala by staphylococcal Alr1 is likely just one mechanism by which *S. aureus* counters the inhibition of Ddl by weak organic acid anions. Additionally, *S. aureus* may also utilize D-Ala produced by the gut microbiota to minimize the impact of acetate intoxication on Ddl (36, 37).

The existence of *pepV* and *dat* within the same operon suggests that these genes may have evolved related functions. In *Lactococcus lactis* the PepV dipeptidase activity was shown to be important for supplying cells with L-Ala which was eventually incorporated into PG (38). In this context, *pepV* and *dat* may have a similar role in modulating the intracellular alanine pool. A surprising finding of our study was that *dat* expression is relatively stable and tightly controlled in *S. aureus* due to its SD motif being located within the coding region of *pepV*. Furthermore, such a genetic arrangement has been linked to translational coupling (39), wherein active translation from the first gene promotes the translation of the following gene in the operon, which in the case of *dat* was not sufficient to overcome acetate toxicity in the *alr1* mutant. Two central mechanisms of translational coupling have been proposed. The first involves secondary and tertiary mRNA structures that either occlude or encompass the SD motif of downstream genes and shield it from ribosomes, thus preventing its translation (40). These mRNA structures can be relieved when a ribosome initiates translation from the first gene of the operon and exposes the downstream intragenic SD sequences to new 30S ribosomal subunits (34). In the second mechanism, continued translation of the first gene of the operon is necessary to increase the abundance of ribosomes in the TIR of the second gene, resulting in its enhanced translation (35). Irrespective of the mechanism of translational coupling, our results suggest that genetic arrangements that promote translational coupling might also limit the overall production of *dat* and thus prevent it from functionally complementing the *alr1* mutant following acetate intoxication. Since our data suggest that Dat primarily promotes flux from D-Ala to D-Glu when Alr1 is active, the tight control of *dat* through translational coupling could prevent the depletion of the intracellular reserves of D-Ala necessary to overcome Ddl inhibition during acetate intoxication. Thus, the elevated D-Ala pool maintained within the cell could represent a strategic adaptation by *S. aureus* to combat Ddl inhibition caused by organic acids typically present in the niches colonized by this bacterium.

In conclusion, our findings demonstrate that Ddl is the primary target of acetate anion intoxication in *S. aureus*. However, other biologically relevant organic anions like lactate, propionate and itaconate could also inhibit the *alr1* mutant similar to acetate. Furthermore, the growth inhibition of the *alr1* mutant by these organic acids could be rescued following D-Ala supplementation, which suggests that Ddl is a *bona fide* and conserved target of various organic acid anions. Indeed, it is tempting to speculate that the robust Alr1 activity leading to the accumulation of millimolar levels of D-Ala may have evolved in part to offset the inhibition of Ddl from the toxic effects of organic anions.

## Acknowledgments

This work was funded by NIH/NIAID R01AI125588 and 2P01A1083211 Metabolomics Core to VCT, 2P01AI083211 Project 4 to TK, respectively. This work was also supported in part by NIH/NIAID R21AI151924 to DRR. X-ray diffraction data were collected at the Life Sciences Collaborative Access Team beamline 21-ID-F at the Advanced Photon Source, Argonne National Laboratory, which is a U.S. Department of Energy (DOE) Office of Science User Facility operated for the DOE Office of Science by Argonne National Laboratory under Contract No. DE-AC02- 06CH11357. Use of the LS-CAT Sector 21 was supported by the Michigan Economic Development Corporation and the Michigan Technology Tri-Corridor (Grant 085P1000817). The University of Nebraska Medical Center Mass Spectrometry and Proteomics Core Facility is administrated through the Office of the Vice Chancellor for Research and supported by state funds from the Nebraska Research Initiative (NRI). Research in the Cava lab is supported by the Swedish Research Council, the Laboratory for Molecular Infection Medicine Sweden (MIMS), Umeå University, the Knut and Alice Wallenberg Foundation (KAW) and the Kempe Foundation. The funders had no role in the study design, data collection, interpretation, and decision to submit this work for publication. The authors have no conflict of interest to declare.

## Data Availability

The atomic coordinates and structure factors have been deposited in the Protein Data Bank, accessible at www.pdb.org, with the PDB ID code 8FFF.

## Materials and Methods

### Bacterial strains and growth conditions

The *S. aureus* WT and mutant strains described in this study were cultured in TSB containing 14 mM glucose. *S. aureus* JE2 mutants were mainly obtained from the Nebraska Transposon Mutant Library (41). These mutants were re-transduced into the WT strain using Ф11- bacteriophage to eliminate any off-target effects. To generate double or triple mutants, the Erm^R^ antibiotic cassette in the transposon mutants was exchanged with Kan^R^ or Tet^R^ cassettes by allelic exchange before introducing an additional mutation. The allelic exchange was performed as described previously (42). In-frame gene deletion mutants were created using a temperature-sensitive vector, pJB38, as described previously (42). *S. aureus* mutants were complemented by inserting the WT allele of mutated genes under the control of their native promoter into the SaPI1 chromosomal site using the pJC1111 suicide vector (43). For experiments involving the over- expression of *ddl* in *S. aureus*, *ddl* was cloned into a CdCl2 inducible multicopy vector, pJB68 (42). The concentration of CdCl2 was titrated to achieve full growth complementation. All bacterial isolates, plasmids, and primers used in this study are listed in Table S5, S6, and S7, respectively.

### Nebraska Transposon Mutant Library (NTML) screen

The NTML mutants were grown in 96-well plates in the presence and absence of 20 mM acetic acid (pH∼6.1) in TSB for 24 hours at 37 °C. The growth of bacteria was determined by measuring the optical density at 600 nm (OD_600_) after 24 hours using a TECAN Infinite 200 spectrophotometer. To account for well-to-well variances that accompany 96-well cultures, the WT strain was independently grown in all the wells of a 96-well plate, both in the presence and absence of acetic acid. Area under the curve (AUC) values for each mutant under a particular condition were obtained by normalizing the values to WT AUC. The graph was generated by plotting the normalized AUC of a mutant under acetate stress versus the control (growth without acetate).

### Competitive fitness assay

The cultures of WT (*S. aureus* JE2) and isogenic mutant strain with either pAQ59 (empty vector) or pAS8 (containing *dat* gene under control of its native promoter) inserted at the SaPI1 chromosomal site were used to assess competitive fitness. Following the growth of these cultures for 24 h, 10^7^ colony forming units (cfu) per milliliter of each strain were used to measure the competitive fitness in presence of 20 mM acetate. The bacterial cfu were enumerated on TSA plates with or without 0.1 mM cadmium chloride immediately after initiation of competition and at 4 h between tested strains allowing the bacteria to undergo approximately seven replications to reach 10^9^ cfu/ ml. The competitive fitness was calculated using the Malthusian parameter for competitors using the following formula: *w* = ln (M*f*/M*i*)/ln (W*f*/W*i*), where *f* and *i* represent cfu counts at final (4 h) and initial (time 0) of competition assay, respectively (11). M and W refer to mutant and WT, respectively.

### Sample collection for mass-spectrometry analysis

Overnight cultures of WT, *alr1* and *dat* mutants were inoculated to an OD_600_ of 0.06 units into 250 ml flasks containing 25ml of TSB 14 mM glucose. Acetic acid (20 mM) was added to the flasks whenever necessary. The flasks were incubated in a shaker incubator at 37 °C and 250 rpm. A total of 10 OD_600_ units of cells were collected following 3 hours of incubation by centrifuging the cultures at 10,000 rpm at 4 °C. The pellet was then washed once with ice-cold saline (0.85% NaCl) and centrifuged again at 10,000 rpm at 4 °C. The bacterial cells were then resuspended in ice cold quenching solution consisting of 60% ethanol, 2 µM Br-ATP and 2 µM ribitol. The cytosolic metabolites were obtained by bead beating the cells, followed by centrifugation. The supernatant was collected and stored at -80 °C until further use. For stable isotope experiments, overnight cultures were inoculated into a chemically defined medium (CDM, (44)) containing ^13^C ^15^N -L-Ala (100 mg/ L) in place of L-Ala and the samples were collected in the exponential phase following 4 hours of incubation at 37 °C.

### Chromatography for mass-spectrometry analysis

The chromatographic separation of PG intermediates was performed by liquid chromatography using XBridge Amide (150 × 2.1 mm ID; 1.7 µm particle size, Waters, USA) analytical column; whereas D-Ala-D-Ala was analysed using XBridge Amide (100 × 2.1 mm ID; 1.7 µm particle size, Waters, USA). A guard XBridge Amide column (20 × 2.1 mm ID; 1.7µm particle size, Waters, USA) was connected in front of the analytical column. Mobile phase A was composed of 10 mM ammonium acetate, 10 mM ammonium hydroxide containing 5 % acetonitrile in LC-MS grade water; mobile phase B was 100% LC-MS grade acetonitrile. The column was maintained at 35 °C and the autosampler temperature was maintained at 5 °C. The gradient was started with the A/B solvent ratio at 15/85 for over 1 minute, followed by a gradual increase of A. A was reduced to 15% after separation and elution of all the interested compounds and equilibrated for 6.0 minutes before the next run. The needle was washed with 1 mL of strong wash solvent containing 100% acetonitrile followed by 1 mL of weak wash solvent comprised of 10% aqueous methanol after each injection. The sample injection volume was 5µl.

Chiral separation of D- and L-isomers of alanine and glutamate was achieved on Astec CHIROBIOTIC^®^ T column (150 x 2.1 mm, 5 µm particle size) from Supelco. Mobile phase A was 20 mM ammonium acetate and mobile phase B was 100% ethanol. The mobile phase composition was 40:60 v/v of A:B in isocratic elution mode pumped at 100 μL/min flow rate. The injection volume was 5 μL and the column was maintained at room temperature. Multiple reaction monitoring (MRM) for D- and L- isomers of alanine are listed in Table S8. All other MS parameters are discussed in the LC-MS/MS analysis section. The L-enantiomer of alanine and glutamate elutes faster than their D-counterparts. The total run time was 15 minutes.

### Targeted LC-MS/MS analysis

Triple-quadrupole-ion trap hybrid mass spectrometer viz., QTRAP 6500+ (Sciex, USA) connected with Waters UPLC was used for targeted analysis. The QTRAP 6500+ was operated in polarity switching mode for targeted quantitation of amino acids through the Multiple Reaction Monitoring (MRM) process. LC-MS MRM data for each metabolite was acquired in centroid mode as a default setting. MRM details for each analyte are listed in Table S8. The optimized electrospray ionization (ESI) parameters were as follows: electrospray ion voltage of -4200 V and 5500 V in negative and positive mode, respectively, source temperature of 500 °C, curtain gas of 40, and gas 1 and 2 of 40 and 40 psi, respectively. Compound-specific parameters were optimized for each compound using manual tuning. These parameters include a declustering potential (DP) of 65 V and -60 V in positive and negative mode, respectively, entrance potential (EP) of 10 V and -10 V in positive and negative mode, respectively, and collision cell exit potential (CXP) maintained at 10 V and -10 V in positive and negative mode respectively. Other compound-specific parameters, such as Q1, Q3, and collision energies, are listed in Table S8. MRM conditions for PG intermediates were adopted from Vemula *et al* (45).

### High Resolution Mass Spectrometry

HRMS Orbitrap (Exploris 480) operated in polarity switching mode was used for the untargeted analysis of isotopologues of D-Ala-D-Ala and D-Glu in data-dependent MS/MS acquisition mode (DDA). Electrospray ionization (ESI) parameters were optimized are as follows: electrospray ion voltage of -2700V and 3500V in negative and positive mode respectively, Ion transfer tube temperature was maintained at 350°C, m/z scan range was 140-180 Da for non-chiral LC-method using Amide column whereas, it was 80-160 Da for chiral column method. Sheath gas, auxiliary gas and sweep gas were optimized according to the UHPLC flow rate. Orbitrap resolution for precursor ion as well as for fragment ion scan was maintained at 240000 and 60000 respectively. Normalized collision energies at 30, 50 and 150% were used for the fragmentation. Data was acquired in profile mode. Xcaliber software from Thermo was used for instrument control and data acquisition. This software was equipped with Qual-, Quant- and FreeStyle browsers which were used for profiling metabolites and their isotopologues in all samples. Selected precursor ion for each isotopologue is listed in Table S9. Identification and detection of all metabolites was aided by the Compound Discoverer (CD) software procured from Thermo USA. The KEGG and HMDB databases plugged-in with CD software were used for metabolite identifications and annotations. Mass accuracy for all the ions was maintained at or below 5 ppm. To correct for natural abundance, we utilized FluxFix, an open-source online software (46), and independently verified these calculations using the ChemCalc software (47).

### Fractional contribution of D-Ala-D-Ala from imported ^13^C ^15^N -L-Ala

An estimate of the fractional contribution (FC) of labeled carbon from ^13^C ^15^N -L-Ala tracer incorporated into the intracellular D-Ala-D-Ala pool was calculated using equation 1, as previously described (48).

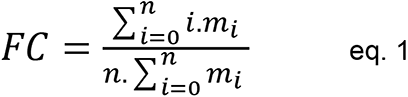

where, *n* is the number of carbon atoms in D-Ala-D-Ala, *i* represents the various carbon isotopologues of D-Ala-D-Ala and *m* the abundance of the D-Ala-D-Ala isotopologues.

### Transcription site identification of the *dat* operon

The adaptor- and radiation-free transcription start site (ARF-TSS) identification method was employed to identify the 5′-UTR region of the *dat* operon (49). In brief, 1 ug of RNA isolated from JE2 WT was subjected to reverse transcription by using 5′-phosphorylated primer pepV_TSS_R1 and the first strand cDNA synthesis kit (Invitrogen, Superscript III First-Strand Synthesis System). RNA was degraded by using 1M NaOH at 65 °C for 30 min and then neutralized with 1M HCl. The resultant cDNA was ligated by using T4 RNA Ligase I (Thermo Scientific) to generate a circular cDNA. Two inverse primers: pepV_TSS_R2 and pepV_TSS_F3 were used to amplify the circular cDNA. The amplified product was cloned into a TOPO Cloning vector and then sequenced using M13F(-20) and M13R primers. All the primers used in this procedure are mentioned in Table S7.

### Quantitative real-time PCR

Quantitative real-time PCR was performed to estimate the transcript levels of *dat*, *ddl* and *murI* in the presence and absence of acetate. The samples were collected during the exponential growth phase and RNA was isolated using a Qiagen RNA isolation kit following the manufacturer’s protocol. A total of 500 ng of RNA was used to synthesize cDNA using the QuantiTech reverse transcription kit (Qiagen). The cDNA samples were then diluted 1:10 and used as a template to perform RT-qPCR. The RT-qPCR was carried out using SYBR green master mix (Roche Applied Science) in a QuantiFast light cycler (Applied Biosystems). The relative transcript levels were estimated by using the comparative threshold cycle method (ΔΔCT) and *sigA* was used as the internal control for normalization. Primers used to perform RT-qPCR are listed in Table S7.

### Muropeptide analysis

The WT and isogenic mutants were inoculated to an OD_600_ of 0.06 into 1-liter flasks containing 100 mL of TSB 14 mM glucose. Acetic acid (20 mM) was added to the media when appropriate. A total of 95 OD_600_ units of cells were collected following 6 hours of growth at 37 °C, 250 rpm. The pelleted cells were then resuspended in 50 % SDS and boiled for 3 hours. Once boiled, cell wall material was pelleted by ultracentrifugation and washed with water. Clean sacculi was digested with muramidase (100 µg/ml) and soluble muropeptides reduced using 0.5 M sodium borate pH 9.5 and 10 mg/mL sodium borohydride. The pH of the samples was then adjusted to 3.5 with phosphoric acid. UPLC analyses was performed on a Waters-UPLC system equipped with an ACQUITY UPLC BEH C18 Column, 130 Å, 1.7 µm, 2.1 mm × 150 mm (Waters Corporation, USA) and identified at Abs. 204 nm. Muropeptides were separated using a linear gradient from buffer A (0.1 % formic acid in water) to buffer B (0.1 % formic acid in acetonitrile). Identification of individual peaks was assigned by comparison of the retention times and profiles to validated chromatograms (50–52). The identity of peak belonging to disaccharide tripeptide, NAG-NAM-AEK (M3) was assigned by mass spectrometry using UPLC system coupled to a Xevo G2/XS Q-TOF mass spectrometer (Waters Corp.). Data acquisition and processing were performed using UNIFI software package (Waters Corp.). The relative amount of each muropeptide was calculated relative to the total area of the chromatogram. Representative chromatograms for each sample type are depicted in (Figure 4-figure supplement 1). The abundance of PG (total PG) was assessed by normalizing the total area of the chromatogram to the OD_600_. The degree of cross-linking refers to the number of peptide bridges and was calculated as % of dimers + % of trimers x 2 + % of tetramers x 3 (53).

### Protein purification

The coding region of *ddl* was cloned into pET28a vector to generate a C-terminal 6×His tag fusion protein before being transferred into *E. coli* BL21(DE3). The cells were grown in Luria Broth Media (Research Product Internationals) containing 50 µg/mL Kanamycin (Gold Biotechnology) at 37 °C. When OD_600_ reached 0.6, 1 mM IPTG (Gold Biotechnology) was added to induce the protein expression. The cells were harvested by centrifugation (3724 *g*) after inducing them at 16 °C for 20 h. The harvested cells were resuspended in lysis buffer comprising 25 mM Tris pH 7.5, 150 mM NaCl, and 5 mM 2-Mercaptoethanol. The cells were lysed by adding Lysozyme (MP-Biomedicals) and DNase I (Roche Applied Sciences) and incubating them on ice for 30 minutes. Then cells were subjected to sonication (Sonicator 3000, Misonix) to further lyse the cells. The crude cell lysate was refined by centrifuging at 18514 *g* for 40 min (Fixed angle rotor, 5810-R Centrifuge, Eppendorf). The clarified lysate was applied to a 5 mL HisTrap™TALON™ crude cobalt column (Cytiva) after equilibrating the column with lysis buffer. The column was washed using the same buffer and the protein was eluted isocratically using 150 mM imidazole-containing buffer. The purified protein was dialyzed in 20 mM Tris pH 8.0 buffer and 0.5 mM Tris (2-carboxyethyl) phosphine to use in crystallization experiments and biochemical assays.

### Crystallization of Ddl and data collection

The crystals of Ddl in complex with acetate were obtained by co-crystallization experiments using the hanging drop vapor diffusion method. The 10 mg/mL of protein was incubated with 30 mM potassium acetate, 5 mM magnesium chloride hexahydrate, and 1 mM ADP for 20 min before the crystallization experiments. The co-crystals were achieved in crystallization drop against a well solution consisting of 0.2 M sodium thiocyanate and 20 % polyethylene glycol monomethyl ether 2000. The crystals were flash cooled in liquid nitrogen immediately after adding 40% polyethylene glycol 3350 to the crystallization drop for cryoprotection. The data were collected at the Advance Photon Source Argonne National Laboratory (APS-ANL, IL), LS-CAT ID-F beamline.

### Ddl enzyme kinetic assays

The Invitrogen™ EnzChek™ Phosphate Assay Kit was used to detect the release of inorganic phosphate by continuously monitoring the absorbance at 360 nm. The reaction components were added as specified by the kit with 200 nm Ddl (containing 1mM MgCl2), 100 mM Potassium chloride, and ATP. The reaction mixture was incubated for 10 min and D-Ala substrate was added to initiate the reaction. The inhibition of Ddl by acetate was determined using various concentrations of sodium acetate, D-Ala, and ATP to determine kinetic parameters.

### Data processing and refinement

The data was processed by CCP4 software (54) and *S. aureus* D-alanyl D-alanine ligase apoprotein (PDB:2I87) was used for the molecular replacement followed by a rigid body refinement using PHENIX (55). Manual model refinement was performed using Coot (56). The XYZ coordinate, B-factor, occupancy, and real space refinements were executed using PHENIX between manual model refinements. The acetate was modeled using eLBOW and positioned at the corresponding difference density. The structure was refined using PHENIX and validated using Molprobity (57).

### Molecular Docking Experiments

The docking experiments of small organic acids were performed with the acetate-bound Ddl structure (PDB:8FFF) with acetate removed. The protein structure was first prepared with the protein preparation wizard. The lactate, propionate and itaconate ligands were prepared by LigPrep. The docking experiments were performed using Schrödinger Glide (New York, NY).

### Differential Scanning Fluorometry

The reaction mixture was prepared using 22 µM Ddl, 5 mM magnesium chloride, 100 mM potassium chloride, 1 mM ADP, 300 mM potassium acetate, and 20 mM Tris pH 7.5 buffer as required. The SyPro orange dye was added to a final concentration of 1 X Protein Thermal Shift™ Dye (Thermofisher) in the reaction mixture. The reactions were performed in triplicate. The samples were centrifuged in MicroAmp™ Optical 96-Well Reaction Plate (Applied Biosystems) at 2325 *g* for 10 minutes. The protein denaturation was monitored by obtaining the fluorescence signal by increasing the temperature from 22 °C - 95 °C at 0.5 °C/minute rate using QuantStudio 3 real-time PCR (ThermoFisher). The melting temperature (Tm) was determined by calculating the derivative of the fluorescent signal and identifying the centroid of the observed melting peak.

## References

1. Passalacqua KD, Charbonneau ME, & O’Riordan MXD (2016) Bacterial metabolism shapes the host-pathogen interface. Microbiol Spectr 4(3).

2. Brestoff JR & Artis D (2013) Commensal bacteria at the interface of host metabolism and the immune system. Nat Immunol 14(7):676–684.

3. Tomlinson KL, et al. (2021) *Staphylococcus aureus* induces an itaconate-dominated immunometabolic response that drives biofilm formation. Nat Commun 12(1):1399.

4. Heim CE, et al. (2020) Lactate production by *Staphylococcus aureus* biofilm inhibits HDAC11 to reprogramme the host immune response during persistent infection. Nat Microbiol 5(10):1271-1284.

5. Schlatterer K, Peschel A, & Kretschmer D (2021) Short-chain fatty acid and FFAR2 Activation - A new option for treating infections? Front Cell Infect Microbiol 11:785833.

6. Salmond CV, Kroll RG, & Booth IR (1984) The effect of food preservatives on pH homeostasis in *Escherichia coli*. J Gen Microbiol 130(11):2845–2850.

7. Roe AJ, McLaggan D, Davidson I, O’Byrne C, & Booth IR (1998) Perturbation of anion balance during inhibition of growth of *Escherichia coli* by weak acids. J Bacteriol 180(4):767–772.

8. Cummings JH, Pomare EW, Branch WJ, Naylor CP, & Macfarlane GT (1987) Short chain fatty acids in human large intestine, portal, hepatic and venous blood. Gut 28(10):1221–1227.

9. Correa-Oliveira R, Fachi JL, Vieira A, Sato FT, & Vinolo MA (2016) Regulation of immune cell function by short-chain fatty acids. Clin Transl Immunology 5(4):e73.

10. Hosmer J, McEwan AG, & Kappler U (2024) Bacterial acetate metabolism and its influence on human epithelia. Emerg Top Life Sci 8(1):1–13.

11. Acton DS, Plat-Sinnige MJ, van Wamel W, de Groot N, & van Belkum A (2009) Intestinal carriage of *Staphylococcus aureus*: how does its frequency compare with that of nasal carriage and what is its clinical impact? Eur J Clin Microbiol Infect Dis 28(2):115–127.

12. Piewngam P, et al. (2023) Probiotic for pathogen-specific *Staphylococcus aureus* decolonisation in Thailand: a phase 2, double-blind, randomised, placebo-controlled trial. Lancet Microbe 4(2):e75–e83.

13. Thomas VC, et al. (2014) A central role for carbon-overflow pathways in the modulation of bacterial cell death. PLoS Pathog 10(6):e1004205.

14. Alqarzaee AA, et al. (2021) Staphylococcal ClpXP protease targets the cellular antioxidant system to eliminate fitness-compromised cells in stationary phase. Proc Natl Acad Sci U S A 118(47).

15. Mordukhova EA & Pan JG (2013) Evolved cobalamin-independent methionine synthase (MetE) improves the acetate and thermal tolerance of *Escherichia coli*. Appl Environ Microbiol 79(24):7905–7915.

16. Roe AJ, O’Byrne C, McLaggan D, & Booth IR (2002) Inhibition of *Escherichia coli* growth by acetic acid: a problem with methionine biosynthesis and homocysteine toxicity. Microbiology (Reading) 148(Pt 7):2215–2222.

17. Gries CM, et al. (2016) Potassium uptake modulates Staphylococcus aureus metabolism. mSphere 1(3).

18. Carpenter CE & Broadbent JR (2009) External concentration of organic acid anions and pH: key independent variables for studying how organic acids inhibit growth of bacteria in mildly acidic foods. J Food Sci 74(1):R12–15.

19. Halsey CR, et al. (2017) Amino acid catabolism in *Staphylococcus aureus* and the function of carbon catabolite repression. mBio 8(1).

20. Moscoso M, Garcia P, Cabral MP, Rumbo C, & Bou G (2018) A D-Alanine auxotrophic live vaccine is effective against lethal infection caused by *Staphylococcus aureus*. Virulence 9(1):604–620.

21. Cava F, Lam H, de Pedro MA, & Waldor MK (2011) Emerging knowledge of regulatory roles of D-amino acids in bacteria. Cell Mol Life Sci 68(5):817–831.

22. Burlak C, et al. (2007) Global analysis of community-associated methicillin-resistant *Staphylococcus aureus* exoproteins reveals molecules produced in vitro and during infection. Cell Microbiol 9(5):1172–1190.

23. Goldstein JM, Kordula T, Moon JL, Mayo JA, & Travis J (2005) Characterization of an extracellular dipeptidase from *Streptococcus gordonii* FSS2. Infect Immun 73(2):1256–1259.

24. Liu S, et al. (2006) Allosteric inhibition of *Staphylococcus aureus* D-Alanine:D-Alanine ligase revealed by crystallographic studies. Proc Natl Acad Sci U S A 103(41):15178–15183.

25. Lu Y, Xu H, & Zhao X (2010) Crystal structure of the apo form of D-Alanine:D-Alanine ligase (Ddl) from *Streptococcus mutans*. Protein Pept Lett 17(8):1053–1057.

26. Pederick JL, Thompson AP, Bell SG, & Bruning JB (2020) D-Alanine-D-alanine ligase as a model for the activation of ATP-grasp enzymes by monovalent cations. J Biol Chem 295(23):7894–7904.

27. Huynh KH, et al. (2015) The crystal structure of the D-alanine-D-alanine ligase from *Acinetobacter baumannii* suggests a flexible conformational change in the central domain before nucleotide binding. J Microbiol 53(11):776–782.

28. Bruning JB, Murillo AC, Chacon O, Barletta RG, & Sacchettini JC (2011) Structure of the *Mycobacterium tuberculosis* D-Alanine:D-Alanine ligase, a target of the antituberculosis drug D-Cycloserine. Antimicrob Agents Chemother 55(1):291–301.

29. Russell JB & Diez-Gonzalez F (1998) The effects of fermentation acids on bacterial growth. Adv Microb Physiol 39:205–234.

30. Russell JB (1992) Another explanation for the toxicity of fermentation acids at low pH: anion accumulation versus uncoupling. Journal of Applied Bacteriology 73(5):363–370.

31. Wolfe AJ (2005) The acetate switch. Microbiol Mol Biol Rev 69(1):12–50.

32. Somerville GA & Proctor RA (2009) At the crossroads of bacterial metabolism and virulence factor synthesis in staphylococci. Microbiol Mol Biol Rev 73(2):233–248.

33. Hammes WP & Neuhaus FC (1974) On the specificity of phospho-N-acetylmuramyl-pentapeptide translocase. The peptide subunit of uridine diphosphate-N-actylmuramyl- pentapeptide. J Biol Chem 249(10):3140–3150.

34. Sobral RG, Ludovice AM, de Lencastre H, & Tomasz A (2006) Role of *murF* in cell wall biosynthesis: isolation and characterization of a murF conditional mutant of *Staphylococcus aureus*. J Bacteriol 188(7):2543–2553.

35. Vemula H, Ayon NJ, & Gutheil WG (2016) Cytoplasmic peptidoglycan intermediate levels in *Staphylococcus aureus*. Biochimie 121:72–78.

36. Matsumoto M, et al. (2018) Free D-amino acids produced by commensal bacteria in the colonic lumen. Sci Rep 8(1):17915.

37. Lee CJ, et al. (2022) Profiling of d-alanine production by the microbial isolates of rat gut microbiota. FASEB J 36(8):e22446.

38. Huang C, Hernandez-Valdes JA, Kuipers OP, & Kok J (2020) Lysis of a *Lactococcus lactis* dipeptidase mutant and rescue by mutation in the pleiotropic regulator CodY. Appl Environ Microbiol 86(8).

39. Huber M, et al. (2019) Translational coupling via termination-reinitiation in archaea and bacteria. Nat Commun 10(1):4006.

40. Rex G, Surin B, Besse G, Schneppe B, & McCarthy JE (1994) The mechanism of translational coupling in Escherichia coli. Higher order structure in the atpHA mRNA acts as a conformational switch regulating the access of de novo initiating ribosomes. J Biol Chem 269(27):18118–18127.

41. Fey PD, et al. (2013) A genetic resource for rapid and comprehensive phenotype screening of nonessential *Staphylococcus aureus* genes. mBio 4(1):e00537–00512.

42. Bose JL, Fey PD, & Bayles KW (2013) Genetic tools to enhance the study of gene function and regulation in *Staphylococcus aureus*. Appl Environ Microbiol 79(7):2218–2224.

43. Chen J, Yoong P, Ram G, Torres VJ, & Novick RP (2014) Single-copy vectors for integration at the SaPI1 attachment site for *Staphylococcus aureus*. Plasmid 76:1–7.

44. Hussain M, Hastings JG, & White PJ (1991) A chemically defined medium for slime production by coagulase-negative staphylococci. J Med Microbiol 34(3):143–147.

45. Vemula H, Bobba S, Putty S, Barbara JE, & Gutheil WG (2014) Ion-pairing liquid chromatography-tandem mass spectrometry-based quantification of uridine diphosphate-linked intermediates in the *Staphylococcus aureus* cell wall biosynthesis pathway. Anal Biochem 465:12–19.

46. Trefely S, Ashwell P, & Snyder NW (2016) FluxFix: automatic isotopologue normalization for metabolic tracer analysis. BMC Bioinformatics 17(1):485.

47. Patiny L & Borel A (2013) ChemCalc: a building block for tomorrow’s chemical infrastructure. J Chem Inf Model 53(5):1223–1228.

48. Fendt SM, et al. (2013) Metformin decreases glucose oxidation and increases the dependency of prostate cancer cells on reductive glutamine metabolism. Cancer Res 73(14):4429–4438.

49. Wang C, Lee J, Deng Y, Tao F, & Zhang LH (2012) ARF-TSS: an alternative method for identification of transcription start site in bacteria. Biotechniques 52(4).

50. de Jonge BL, Chang YS, Gage D, & Tomasz A (1992) Peptidoglycan composition of a highly methicillin-resistant Staphylococcus aureus strain. The role of penicillin binding protein 2A. J Biol Chem 267(16):11248–11254.

51. De Jonge BL, Gage D, & Xu N (2002) The carboxyl terminus of peptidoglycan stem peptides is a determinant for methicillin resistance in *Staphylococcus aureus*. Antimicrob Agents Chemother 46(10):3151–3155.

52. Kuhner D, Stahl M, Demircioglu DD, & Bertsche U (2014) From cells to muropeptide structures in 24 h: peptidoglycan mapping by UPLC-MS. Sci Rep 4:7494.

53. Alvarez L, Hernandez SB, de Pedro MA, & Cava F (2016) Ultra-sensitive, high-resolution liquid chromatography methods for the high-throughput quantitative analysis of bacterial cell wall chemistry and structure. Methods Mol Biol 1440:11–27.

54. Winn MD, et al. (2011) Overview of the CCP4 suite and current developments. Acta Crystallogr D Biol Crystallogr 67(Pt 4):235–242.

55. Afonine PV, et al. (2012) Towards automated crystallographic structure refinement with phenix.refine. Acta Crystallogr D Biol Crystallogr 68(Pt 4):352–367.

56. Emsley P, Lohkamp B, Scott WG, & Cowtan K (2010) Features and development of Coot. Acta Crystallogr D Biol Crystallogr 66(Pt 4):486–501.

57. Chen VB, et al. (2010) MolProbity: all-atom structure validation for macromolecular crystallography. Acta Crystallogr D Biol Crystallogr 66(Pt 1):12–21.

58. Lambert RJ & Pearson J (2000) Susceptibility testing: accurate and reproducible minimum inhibitory concentration (MIC) and non-inhibitory concentration (NIC) values. J Appl Microbiol 88(5):784–790.

59. Kreiswirth BN, et al. (1983) The toxic shock syndrome exotoxin structural gene is not detectably transmitted by a prophage. Nature 305(5936):709–712.

60. Lee CY, Buranen SL, & Ye ZH (1991) Construction of single-copy integration vectors for *Staphylococcus aureus*. Gene 103(1):101–105.

61. Jacquet R, et al. (2019) Dual gene expression analysis identifies factors associated with *Staphylococcus aureus* virulence in diabetic mice. Infect Immun 87(5):e00163–00119.

