## Supplementary Data for "*Staphylococcus aureus* counters organic acid anion-mediated inhibition of peptidoglycan cross-linking through robust alanine racemase activity"

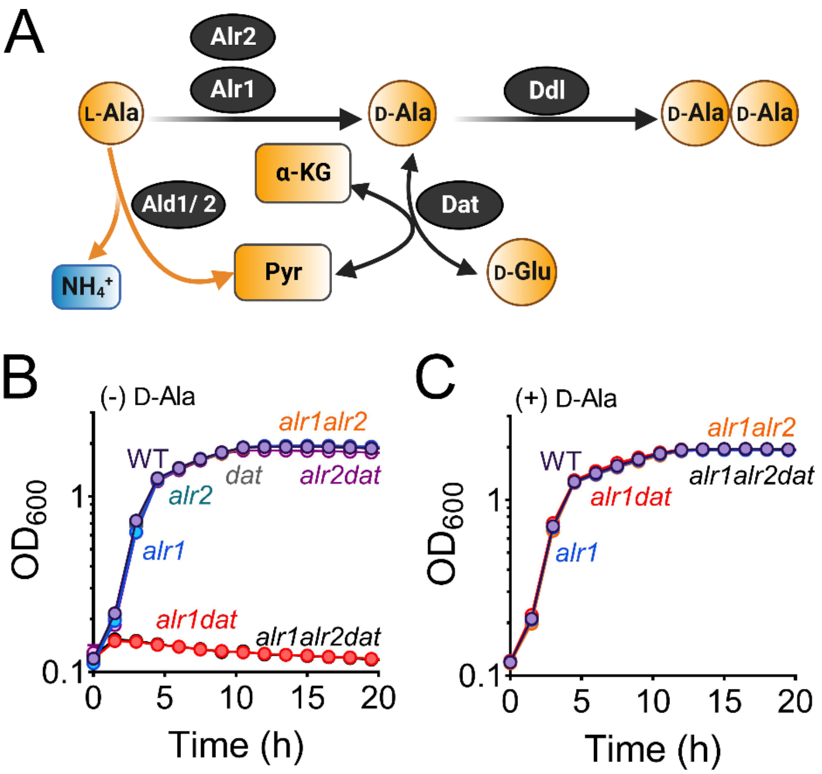


**Figure 1-figure supplement 1. Alr1 and Dat are the primary routes of D-Ala production in *S. aureus.* (A)** Schematic of the predicted D-Ala-D-Ala generating pathways in *S. aureus*. Growth curves of various *S. aureus* strains grown in the **(B)** absence or **(C)** presence of 5 mM D-Ala (n=3, mean ± SD).

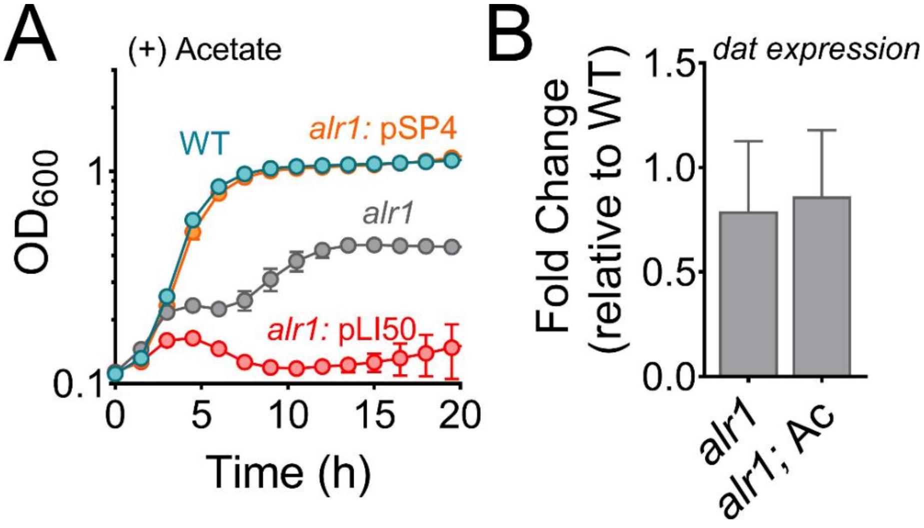


**Figure 2-figure supplement 1. Overexpression of *dat* rescues the growth defect of the *alr1* mutant (A)** *dat* was cloned in a multicopy vector (pSP4) controlled by its native promoter. The *S. aureus* strains containing pSP4 and pLI50 (empty vector) were grown in TSB supplemented with 20 mM acetate **(B)** RT-qPCR analysis of *dat* expression in the *alr1* mutant in the presence or absence of 20 mM acetic acid (n=3, mean ± SD).

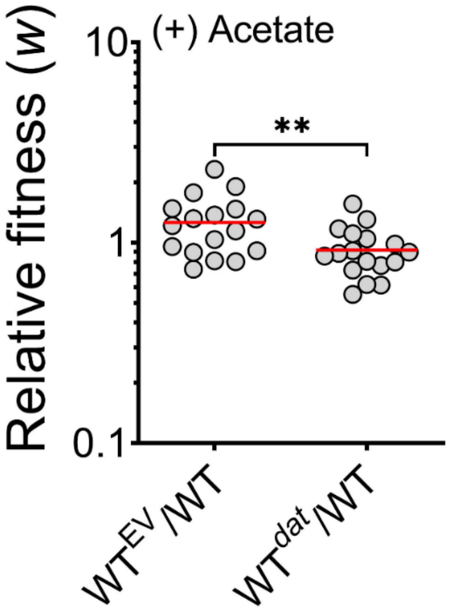


**Figure 3-figure supplement 1. Increased *dat* expression causes a fitness defect.** The mean competitive fitness (*w*) was determined by co-culturing the WT strain with an isogenic mutant that contained either the empty pAQ59 vector integrated into the SaPI chromosomal site ($WT^{EV}$) or the pAS8 vector containing *dat* under the control of its native promoter ($WT^{dat}$) (n=18, the dotted lines indicate the median and quartiles). **, P value <0.01.

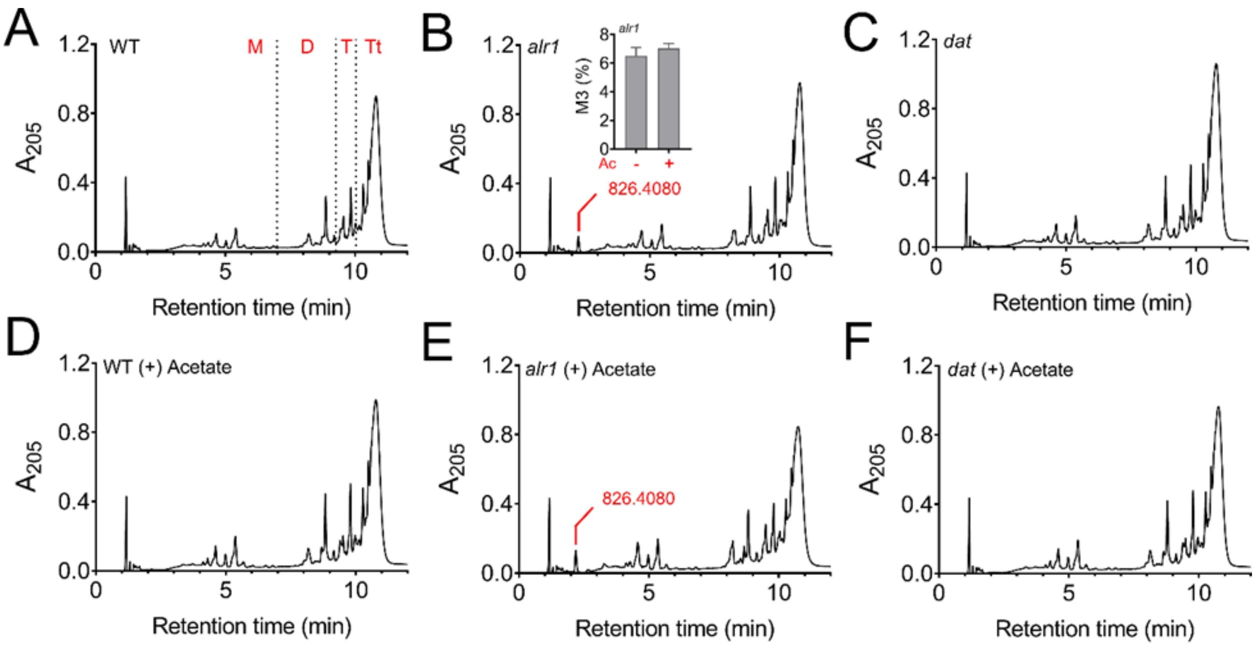


**Figure 4-figure supplement 1. Muropeptide analysis.** Representative **c**hromatograms of muropeptide extracts from **(A-C)** WT, *alr1* and *dat* mutants, and following **(D-F)** acetate intoxication. A unique peak corresponding to NAG-NAM-AEK (M3, m/z, Da: 826.4080) was identified in the *alr1* mutant. The peak area of M3 was normalized to the total area of peaks observed in the chromatogram and expressed as percent (see inset figure in **B**, n=3, mean ± SD). M, monomer; D, dimer; T, trimer, Tt, tetramer.

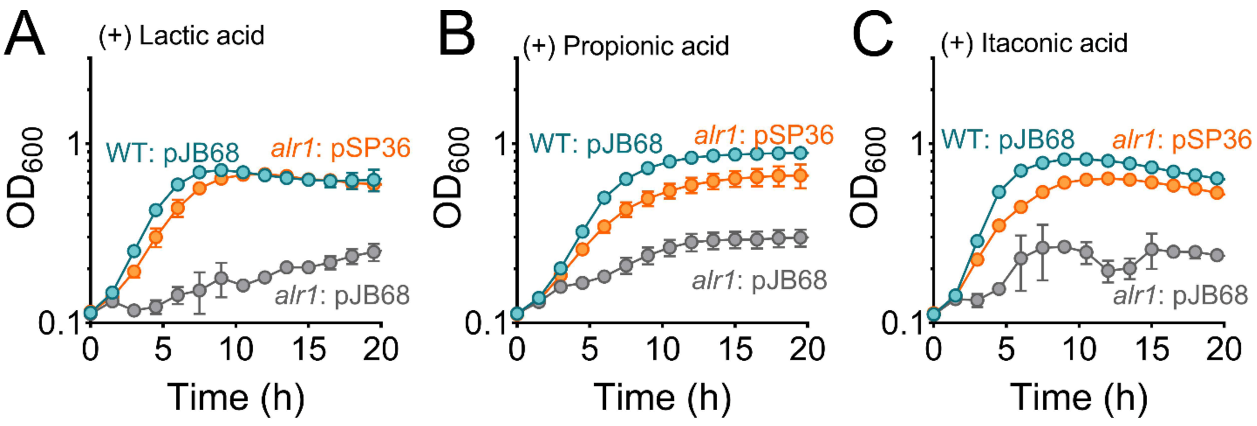


**Figure 6-figure supplement 1. Overexpression of *ddl* rescues the growth defect of the *alr1* mutant** The growth ($OD_{600}$) of the WT and *alr1* mutants overexpressing Ddl (pSP36; cadmium inducible expression of *ddl*) in TSB supplemented with **(A)** lactic acid (40 mM), **(B)** propionic acid (20 mM) and **(C)** Itaconic acid (20 mM) in the presence or absence of 5 mM D-Ala (n=3, mean ± SD).

**Table S1: Effect of acetate on Ddl activity**

| Substrate | Condition | $K_m$ (mM) | $V_{max}$ (µM $min^{-1}$) | $k_{cat}$ ($min^{-1}$) |
|---|---|---|---|---|
| D-**Ala** | 0 mM Acetate | 7.0 ± 0.6 | 16.0 ± 0.4 | 80.1 ± 2.1 |
| | 100 mM Acetate | 6.8 ± 0.6 | 10.9 ± 0.3 | 54.4 ± 1.6 |
| | 300 mM Acetate | 8.4 ± 0.6 | 6.3 ± 0.1 | 31.5 ± 0.7 |
| **ATP** | 0 mM Acetate | 0.6 ± 0.1 | 12.3 ± 1.1 | 61.7 ± 5.7 |
| | 200 mM Acetate | 0.8 ± 0.2 | 6.9 ± 0.8 | 34.4 ± 3.8 |
| | 300 mM Acetate | 0.8 ± 0.3 | 4.0 ± 0.7 | 19.9 ± 3.6 |

**Table S2: Impact of acetate on Ddl stability assessed by DSF**

| Sample | Tm D (ºC) | ΔTm D (ºC)* |
|---|---|---|
| Ddl | 45.0 ± 0.0 | - |
| Ddl + Sodium acetate (Ac) | 48.7 ± 0.0 | 3.7 |
| Ddl + ATP | 46.4 ± 0.0 | 1.4 |
| Ddl + ATP + Ac | 48.9 ± 0.1 | 3.9 |
| Ddl + ADP | 41.9 ± 0.1 | -3.2 |
| Ddl + ADP + Ac | 49.7 ± 0.1 | 4.0 |
| Ddl + D-Ala | 49.2 ± 0.1 | 4.2 |
| Ddl + D-Ala + Ac | 49.6 ± 0.0 | 4.5 |
| Ddl + D-Ala + ATP + Ac | 47.1 ± 0.1 | 2.0 |
| Ddl + D-Ala + ADP + Ac | 49.8 ± 0.0 | 4.8 |

*The ΔTm D values are calculated as the difference in melting temperature of the Ddl apo protein to Ddl with added substrates or acetate inhibitor.

**Table S3: Refinement statistics of Ddl/Acetate structure**

| | |
|---|---|
| Resolution Range | 48.21 – 1.92 (1.989 – 1.92) |
| Space group | P 2 21 21 |
| Unit cell | 55.123 65.817 99.423 90 90 90 |
| Total reflections | 173526 (12202) |
| Unique reflections | 20366 (2806) |
| Multiplicity | 8.5 (9.3) |
| Completeness (%) | 94.80 (50.46) |
| Mean I/sigma(I) | 10.99 (1.72) |
| Wilson B-factor | 37.29 |
| R-merge | 0.097 (0.33) |
| R-meas | 0.104 (0.35) |
| R-pim | 0.036 (0.12) |
| CC1/2 | 0.998 (0.95) |
| CC* | 0.999 (0.99) |
| Reflections used in refinement | 26870 (1416) |
| Reflections used for R-free | 1601 (92) |
| R-work | 0.21 (0.42) |
| R-free | 0.27 (0.47) |
| CC (work) | 0.223 (0.07) |
| CC (free) | 0.217 (-0.16) |
| Number of non-hydrogen atoms | 2963 |
| macromolecules | 2777 |
| ligands | 10 |
| solvent | 176 |
| Protein residues | 355 |
| RMS(bonds) | 0.009 |
| RMS(angles) | 1.10 |
| Ramachandran favored (%) | 93.70 |
| Ramachandran allowed (%) | 6.02 |
| Ramachandran outliers (%) | 0.29 |
| Rotamer outliers (%) | 0.00 |

| | |
|---|---|
| Clashscore | 6.88 |
| Average B-factor | 43.60 |
| macromolecules | 43.60 |
| ligands | 45.09 |
| solvent | 43.64 |

**Table S4: Glide scores from molecular docking studies of organic anions**

| Organic Anion | ATP binding site | D-Ala binding site |
|---|---|---|
| L-lactate | -4.851 | -4.431 |
| Propionate | -2.317 | -1.883 |
| Itaconate | -3.572 | -3.575 |

**Table S5: Strains used in this study**

| Strains | Description | Source |
|---|---|---|
| *E. coli* Electro-Ten-Blue | General plasmid maintenance strain | Stratagene |
| *S. aureus* RN4220 | Restriction-deficient strain is routinely used as a transformation intermediate | (59) |
| *S. aureus* RN4220: pRN7023 | Restriction deficient strain carrying pRN7023 plasmid containing integrase gene routinely used as a transformation intermediate | (43) |
| *E. coli* DH5α | General plasmid maintenance strain | Thermo Fisher |
| *E. coli* BL21(DE3) | Protein over-expression and purification strain | Novagen |
| *S. aureus* JE2 | *S. aureus* USA300 LAC cured of all 3 native plasmids | (41) |
| JE2 *alr1* | *bursa aurealis* transposon mutant, $Erm^R$ | NTML |
| JE2 *alr1::alr1* | WT copy of *alr1* complemented at the SaPI1 site of *bursa aurealis* transposon mutant, $Erm^R$ | This study |
| JE2 *citZ* | *bursa aurealis* transposon mutant, $Erm^R$ | NTML |
| JE2 *citZalr1* | *bursa aurealis* transposon mutant, $Erm^R$, $Kan^R$ | This study |
| JE2 *dat* | *bursa aurealis* transposon mutant, $Erm^R$ | NTML |
| JE2 Δ*alr2* | Inframe isogenic deletion mutant of JE2 | This study |
| JE2 *alr1*Δ*alr2* | *bursa aurealis* transposon mutant, $Erm^R$, transduced into inframe isogenic deletion mutant JE2 Δ*alr2* | This study |
| JE2 Δ*alr2dat* | *bursa aurealis* transposon mutant, $Erm^R$, transduced into inframe isogenic deletion mutant JE2 Δ*alr2* | This study |
| JE2 *alr1*Δ*alr2dat* | *bursa aurealis* transposon mutant, $Tet^R$, transduced into inframe isogenic deletion mutant JE2 alr1Δ*alr2* | This study |
| JE2 *alr1dat* | *bursa aurealis* transposon mutant, $Erm^R$, $Tet^R$ | This study |
| JE2 alr1: *dat* | pLI50 *dat* plasmid (pSP4) transduced into *bursa aurealis* transposon mutant, $Erm^R$ | |
| JE2 $pepV^{\Delta SD1\text{-}467}$ | Isogenic deletion mutant of SD1 and *pepV* | This study |
| JE2 $alr1pepV^{\Delta SD1\text{-}467}$ | *bursa aurealis* transposon mutant, $Erm^R$ transduced into isogenic deletion mutant of SD1 and *pepV* | This study |
| JE2 Δ*pepV* | Inframe isogenic deletion mutant of JE2 | This study |
| JE2 *alr1*Δ*pepV* | *bursa aurealis* transposon mutant, $Erm^R$ transduced into inframe isogenic deletion mutant JE2 Δ*pepV* | This study |

| | | |
|---|---|---|
| JE2 *pepV*$^{Q12STOP}$ | Glutamine to STOP codon substitution at 12$^{th}$ amino acid position in PepV | This study |
| JE2 *alr1pepV*$^{Q12STOP}$ | *Bursa aurealis* transposon mutant, Erm$^{R}$ transduced into glutamine to STOP codon substitution at 12$^{th}$ amino acid position in PepV | This study |
| JE2 WT: pJB68 | pJB68 transduced into WT JE2 | This study |
| JE2 WT: pSP36 | pSP36 transduced into WT JE2 | This study |
| JE2 *alr1*: pJB68 | pJB68 transduced into JE2 *alr1* | This study |
| JE2 *alr1*: pSP36 | pSP36 transduced into WT *alr1* | This study |
| JE2 WT::pAQ59 | pAQ59 empty vector inserted at the SaPI1 site of WT JE2 | This study |
| JE2 WT::pAS8 | *dat* (under its native promoter) inserted at the SaPI1 site of WT JE2 | This study |

**Table S6: Plasmids used in this study**

| Plasmids | Description | Source |
|---|---|---|
| pLI50 | *E. coli*- *S. aureus* shuttle vector | (60) |
| pJB38 | *E. coli*- *S. aureus* allelic exchange vector | (42) |
| pJC1111 | *E. coli*- *S. aureus* SaPI1 site integration vector | (43) |
| pJB68 | *E. coli*- *S. aureus* cadmium inducible shuttle vector | (42) |
| pET28a | Expression vector for purification of protein in *E. coli* BL21 (DE3) | Novagen |
| pAS3 | pJC1111 based vector for integration of WT copy of *alr1* at the SaPI1 site | This study |
| pAS2 | pJB38 based vector for *alr2* chromosomal deletion | This study |
| pSP4 | pLI50 *dat* (under control of its native promoter) | This study |
| pSP19 | pJB38 based vector for *pepV* SD1 chromosomal deletion | This study |
| pSP20 | pJB38 based vector for *pepV* SD1-467 chromosomal deletion | This study |
| pSP16 | pJB38 based vector for *pepV* chromosomal deletion | This study |
| pSP15 | pJB38 based vector for substitution of chromosomal *pepV* with *pepV*$^{Q12STOP}$ | This study |
| pSP36 | pJB68 based vector for overexpression of *ddl* | This study |
| pSP32 | pET28a based vector for purification of full length Ddl (C-terminal his tag) | This study |
| pAQ59 | *E. coli*- *S. aureus* SaPI1 site integration vector with pSC101 ori region | (61) |
| pAS8 | pAQ59 based vector for integration of *dat* gene under its native promoter at the SaPI1 site | This study |

**Table S7: Primers used in this study**

| Gene/Modification | Primer name | Primer sequence (5' – 3') |
|---|---|---|
| *alr1* | alr1_F | TGCTGACGAACCAGGAGATA |
| | alr1_R | TGTAGTTGGGTCAGTAGCTG |
| *alr1* complementation | alr1_comp_F | CGGCCGCTGCATGCCTGCAGACATGAGCAACGTAAA ATTG |
| | alr1_comp_R | AGCTCGGTACCCGGGGATCCAATGACCTTTAATTACT CTAATGATAAC |
| *citZ* | 1641_F | CAGCGGAGACTAAAATAAGTTC |
| | 1641_R | CCCAATCTCAGATAACATCGTC |
| *dat* | dat_F | ACTATAGGTGGCGGTACTTA |
| | dat_R | ACCATCGGATATCTTCAACG |
| *alr2* deletion | alr2_UP_F | CGAGGCCCTTTCGTCTTCAATACTTAGAAGGTAATGG CTC |
| | alr2_UP2_R | TCATAGCACTTGCTGTCAATGTATTACAC |
| | alr2_DN2_F | ATTGACAGCAAGTGCTATGAATCATGATTC |
| | alr2_DN_R | TTGCATGCCTGCAGGTCGACGCTTCTTCATTTCTATTA ACAAG |
| *dat* complementation | dat promoter_F | CCTTTCGTCTTCAAGAATTCGATGTGAGTAGGACAGA AATG |
| | dat promoter_R | TTTTTTCCATTCGAAATCGACTTCCTTTTTTC |
| | dat_pLI50_F | TCGATTTCGAATGGAAAAAATTTTTTTAAATGGTG |
| | dat_pLI50_R | TTGCATGCCTGCAGGTCGACCGAAAGTTGATAAATTT AAGTAATTTAATC |
| TSS identification of *dat* operon | pepV_TSS_R1 | P-CCATCTCTATGTGCAATTTC |

| | | |
|---|---|---|
| | pepV_TSS_R2 | GCGTCTTCTGATGCTTTTGC |
| | pepV_TSS_F3 | GTCCTCGTAAGGCATTAGAC |
| | M13F (-20) | GTAAAACGACGGCCAG |
| | M13R | CAGGAAACAGCTATGAC |
| *pepV*ΔSD1-467 | RBS1pepV_UP_F | CCTTTCGTCTTCAAGAATTCAGCGACGCAATTAGGAA C |
| | RBS1pepV_UP_R | TTATTCCTCCTTTTTCTATAAGTTAAATTCTATTTTACAT GAAAAG |
| | RBS1pepV_DN_F | TATAGAAAAAGGAGGAATAATATATGGAAAAAATTTTT TTAAATG |
| | RBS1pepV_DN_R | TATAGAAAAAGGAGGAATAATATATGGAAAAAATTTTT TTAAATG |
| *pepV* deletion | pepV_UP2_F | CCTTTCGTCTTCAAGAATTCAACAATTAAAGAAGTAAA AACAAATC |
| | pepV_UP2_R | TTTTTTCCATTCGAAATCGACTTCCTTTTTTC |
| | pepV_DN2_F | TCGATTTCGAATGGAAAAAATTTTTTTAAATGGTG |
| | pepV_DN2_R | TTGCATGCCTGCAGGTCGACTTTCAACTGAAAATGAG AAAC |
| *pepV*Q12STOP | pepV_STOP_UP _F | CCTTTCGTCTTCAAGAATTCCAAATCCGAAAGAATATG C |
| | pepV_STOP_UP _R | TAATGATTTAATCTTCGTATTGTTGAACTTTTTC |
| | pepV_STOP_D N_F | ATACGAAGATTAAATCATTAATGACTTAAAAGGATTATT AG |
| | pepV_STOP_D N_R | TTGCATGCCTGCAGGTCGACAAAGACCTGCGTTTTCA TTATC |
| *ddl* overexpression plasmid | ddl_pJB68_F | TTTATAAGGAGGAAAAACATATGACAAAAGAAAATATT TGTATCG |
| | ddl_pJB68_R | GAATAGGCGCGCCTGAATTCATCCATGATTGAATTTG CTTTAATG |

| | | |
|---|---|---|
| Ddl purification plasmid | ddl_C_Histag_F | CTTTAAGAAGGAGATATACCATGACAAAAGAAAATATTTGTATCG |
| | ddl_C_Histag_R | CAGTGGTGGTGGTGGTGGTGGTCAATTTTGTATTTATTTTTCTGTTTATC |
| *ddl* RT-qPCR | ddl_RT_F | GGGCTTTTTGAAGTTTTGGA |
| | ddl_RT_R | TGGTAACCCTCGATGTTCAA |
| *murF* RT-qPCR | murF_RT_F | TCACAATTGATTCACGAGCA |
| | murF_RT_R | CCCAGCACCATCTTGTAATG |
| *dat* RT-qPCR | dat RT_F | GATGGTTACGTTGCGACATT |
| | dat RT_R | CACCTCGATGTTGAATTGCT |
| *sigA* RT-qPCR | JE2_RT_sigA_F | AACTGAATCCAAGTGATCTTAGTG |
| | JE2_RT_sigA_R | TCATCACCTTGTTCAATACGTTTG |
| *dat* insertion at the SaPI1 site (pAS8) | dat_UP2_F | GAGCCGCTGCATGCCTGCAGGATGTGAGTAGGACAGAAATG |
| | dat_UP_R | TTTTTTCCATTCGAAATCGACTTCCTTTTTTC |
| | dat_DN_F | TCGATTTCGAATGGAAAAAATTTTTTTAAATGGTG |
| | dat_DN_R | AGCTCGGTACCCGGGGATCCCGAAAGTTGATAAATTTAAGTAATTTAATC |

**Table S8: Table of Multiple Reaction Monitoring (MRM) transitions**

| Metabolite | Polarity | MRM (Q1/Q3) | CE (V) | DP (V) | RT | Column |
|---|---|---|---|---|---|---|
| L-Ala | (+) | 90.1 / 44.0 | 17 | 65 | 6.4 | C |
| D-Ala | (+) | 90.1 / 44.0 | 17 | 65 | 9.5 | C |
| D-Ala-D-Ala | (+) | 161.0 / 44.2 | 30.5 | 65 | 3.4 | XB |
| UDP-NAG | (-) | 606.0 / 79.0 | -149 | -80 | 12.7 | XB |
| UDP-NAM | (-) | 678.1 / 79.0 | -120 | -80 | 13.2 | XB |
| UDP-NAM-A | (-) | 749.1 / 403.0 | -42 | -90 | 13.4 | XB |
| UDP-NAM-AE | (-) | 878.2 / 403.0 | -48 | -105 | 14.3 | XB |
| UDP-NAM-AEK | (-) | 1006.2 / 403.0 | -50 | -130 | 14.9 | XB |
| UDP-NAM-AEKAA | (-) | 1148.5 / 403.0 | -55 | -140 | 14.6 | XB |
| NAM | (-) | 292.0 / 89.0 | -16 | -30 | 6.2 | XB |
| NAG | (+) | 204.0 / 138.1 | 18.9 | 30 | 5.6 | XB |
| Br-ATP (IS) | (-) | 588.0 / 159.0 | -38.9 | -60 | 6.0 | XB |
| Ribitol (IS) | (-) | 151.1 / 89.0 | -14.8 | -60 | 5.3 | XB |

CE: Collision energy ;DP: Declustering potential; RT: Retention Time; C: CHIROBIOTIC® T column; XB: XBridge Amide column

**Table S9: HRMS base peak identification of isotopologues**

| | Metabolite | Isotopologues($^{13}C^{15}N$) | Base peak (m/z) |
|---|---|---|---|
| **D-Ala-D-Ala (Positive mode)** | | | |
| 1 | C6H12N2O3 | $C_0N_0$ | 161.0921 |
| 2 | [13]C1C5H12N2O3 | $C_1N_0$ | 162.0954 |
| 3 | [13]C2C4H12N2O3 | $C_2N_0$ | 163.0988 |
| 4 | [13]C3C3H12N2O3 | $C_3N_0$ | 164.1021 |
| 5 | [13]C3C3H12[15]N1N1O3 | $C_3N_1$ | 165.09917 |
| 6 | [13]C3C3H12[15]N2O3 | $C_3N_2$ | 166.0962 |
| 7 | [13]C4C2H12N2O3 | $C_4N_0$ | 165.10549 |
| 8 | [13]C5C1H12N2O3 | $C_5N_0$ | 166.10884 |
| 9 | [13]C6H12N2O3 | $C_6N_0$ | 167.1122 |
| 10 | [13]C6H12[15]N1N1O3 | $C_6N_1$ | 168.1092 |
| 11 | [13]C6H12[15]N2O3 | $C_6N_2$ | 169.1063 |
| 12 | [13]C1C5H12[15]N1N1O3 | $C_1N_1$ | 163.09246 |
| 13 | [13]C1C5H12[15]N2O3 | $C_1N_2$ | 164.0895 |
| 14 | [13]C2C4H12[15]N1N1O3 | $C_2N_1$ | 164.0958 |
| 15 | [13]C2C4H12[15]N2O3 | $C_2N_2$ | 165.09285 |
| 16 | [13]C4C2H12[15]N1N1O3 | $C_4N_1$ | 166.1025 |
| 17 | [13]C4C2H12[15]N2O3 | $C_4N_2$ | 167.09956 |
| 18 | [13]C5C1H12[15]N1N1O3 | $C_5N_1$ | 167.10588 |
| 19 | [13]C5C1H12[15]N2O3 | $C_5N_2$ | 168.1029 |
| 20 | C6H12[15]N1N1O3 | $C_0N_1$ | 162.0891 |
| 21 | C6H12[15]N2O3 | $C_0N_2$ | 163.0861 |
| **D-Glu (Negative mode)** | | | |
| 1 | C5H9NO4 | $C_0N_0$ | 146.04588 |
| 2 | C5H9[15]NO4 | $C_0N_1$ | 147.04292 |
| 3 | [13]C1C4H9NO4 | $C_1N_0$ | 147.04924 |
| 4 | [13]C2C3H9NO4 | $C_2N_0$ | 148.05259 |
| 5 | [13]C2C3H9[15]NO4 | $C_2N_1$ | 149.04963 |
| 6 | [13]C1C4H9[15]NO4 | $C_1N_1$ | 148.04627 |
| 7 | [13]C3C2H9[15]NO4 | $C_3N_1$ | 150.05298 |
| 8 | [13]C4C1H9[15]NO4 | $C_4N_1$ | 151.05634 |
| 9 | [13]C5H9[15]NO4 | $C_5N_1$ | 152.05969 |
| 10 | [13]C3C2H9NO4 | $C_3N_0$ | 149.05595 |
| 11 | [13]C4C1H9NO4 | $C_4N_0$ | 150.0593 |
| 12 | [13]C5H9NO4 | $C_5N_0$ | 151.06266 |

RT: D-Ala-D-Ala, 3.4 mins on 10 cm XBridge amide column; D-Glu, 6.4 mins on CHIROBIOTIC® T column
